# Human brain solute transport quantified by glymphatic MRI-informed biophysics during sleep and sleep deprivation

**DOI:** 10.1101/2023.01.01.522190

**Authors:** Vegard Vinje, Bastian Zapf, Geir Ringstad, Per Kristian Eide, Marie E. Rognes, Kent-Andre Mardal

## Abstract

Whether you are reading, running or sleeping, your brain and its fluid environment continuously interacts to distribute nutrients and clear metabolic waste. Yet, the precise mechanisms for solute transport within the human brain have remained hard to quantify using imaging techniques alone. From multi-modal human brain MRI data sets in sleeping and sleep-deprived subjects, we identify and quantify CSF tracer transport parameters using forward and inverse subject-specific computational modelling. Our findings support the notion that extracellular diffusion alone is not sufficient as a brain-wide tracer transport mechanism. Instead, we show that human MRI observations align well with transport by either substantially enhanced (3.5×) extracellular diffusion in combination with local clearance rates corresponding to a tracer half-life of up to 5 hours, or by extracellular diffusion augmented by advection with brain-wide average flow speeds on the order of 1–9 *µ*m/min. Reduced advection fully explains reduced tracer clearance after sleep-deprivation, supporting the role of sleep and sleep deprivation on human brain clearance.

## Introduction

The brain and its fluid surroundings form a singular environment for solute influx, exchange and clearance, marked by intertwined vascular and extravascular pathways^1, 2^. Indeed, the privileged absence of lymphatic vessels within the brain parenchyma^3^ accentuates other potential modes of metabolic solute transport such as extracellular diffusion^4, 5^, advection by cerebrospinal or interstitial fluid flow^6, 7^, and local clearance across the blood-brain barrier^1^. The introduction of the glymphatic theory^8^ marked the beginning of a resurgence of research into these mechanisms, and their implication in neurodegenerative disease^1, 9^, neurological disorders^10^, stroke^11^, edema^12^, oncology^13^, drug delivery^14^, and sleep^15–17^. Yet, their contribution and relative roles remain under active debate^18–21^, in part due to the lack of direct in-vivo measurements, and proxies offered by diffusion tensor imaging (DTI), contrast-based magnetic resonance imaging (MRI) or fluorescence microscopy.

In the extracellular space (ECS) and across species, diffusion parameters are well-established via experimental, clinical, as well as computational techniques^4, 22^. Moreover, extracellular diffusion appears to dominate advection by interstitial fluid (ISF) flow in the nanoscale ECS^19, 23, 24^. On the other hand, cerebrospinal fluid (CSF) velocities are observed at the order of cm/s in human^25^, and reach tens of *µ*m/s in mice pial perivascular spaces (PVSs)^26^. Whether substantial fluid velocities also manifest in parenchymal PVSs and across species (importantly including in humans) are open questions; notably juxtaposed by experimental observations of bulk ISF rates of the order *µ*m/min in rats^4, 6, 27^, and MRI-guided computational models revealing effective interstitial diffusivities enhanced by 10–25 × in mice^28^.

In humans, CSF tracer (gadobutrol) enrichment after intrathecal injection is characterized by fast transport in the CSF over the first few hours, a brain-wide enrichment over the first 24 hours, followed by decline from 24–48 hours, and with no evidence of tracer remaining in the brain after 4 weeks^10, 29–31^. Intriguingly, the tracer enrichment patterns differ between sleeping and sleep-deprived subjects, both in the cerebral cortex and in the subcortical white matter^17^. Altered brain tracer enrichment also accompanies chronic poor sleep quality^32^. These observations thus complement previous striking reports of the effect of sleep on brain solute influx and clearance in mice^15^. Further, contrast MRI-informed biophysics models reveal that the tracer spreads faster also within the human brain than by extracellular diffusion alone^33, 34^, albeit without pin-pointing or quantifying alternative transport parameters.

*Forward* computational models of macroscale solute transport^33^, diffusion and flow in the ECS^23, 24, 35^, and notably perivascular fluid flow and transport^36–42^ are now effective complementary tools for evaluating physiological hypotheses. Yet, there is an untapped technological potential for *inverse* computational modelling in which biophysics-based models of solute transport are synthesized with multi-modal data to systematically identify and quantify underlying transport parameters. Valnes et al^34^ estimate an effective isotropic diffusion coefficient based on MRI data in a limited set of (three) human subjects, but do not account for other mechanisms. In rats, and also leveraging dynamic contrast-enhanced MRI, Tannenbaum and colleagues have developed and refined an optimal control approach for identifying glymphatic vector fields^43–45^, but without quantifying absolute velocity magnitudes, water influx or local clearance rates.

In this study, we identify and quantify extravascular solute transport parameters in the human brain by combining high-fidelity inverse biophysical modelling with multi-modal MRI data from 24 subjects over 48 hours, including seven subjects deprived of sleep. We represent a comprehensive set of glymphatic-type transport mechanisms including 1) extracellular diffusion, 2) *α*-enhanced extracellular diffusion, 3) *α*-enhanced extracellular diffusion combined with local solute clearance, and 4) extracellular diffusion combined with advection by tissue fluid flow. For the latter two scenarios, we estimate *α* and the local clearance rates *r*, and the fluid flow velocities *φ*, respectively. Our findings (Fig. 1) demonstrate that clinically-observed tracer influx and clearance patterns are compatible with (a) enhanced effective extracellular diffusion with *α* ≈ 3.5 combined with a local clearance rate of *r* ≈ 3.1 ×10^*−*3^/min, or (b) extracellular diffusion augmented by advection with average fluid flow speeds of |*φ*| ≈ 1–8 *µ*m/min. The average fluid flow speeds were reduced by a factor of two during the clearance phase (24–48 hours) in sleep-deprived subjects compared to sleeping reference subjects.

**Figure 1.**
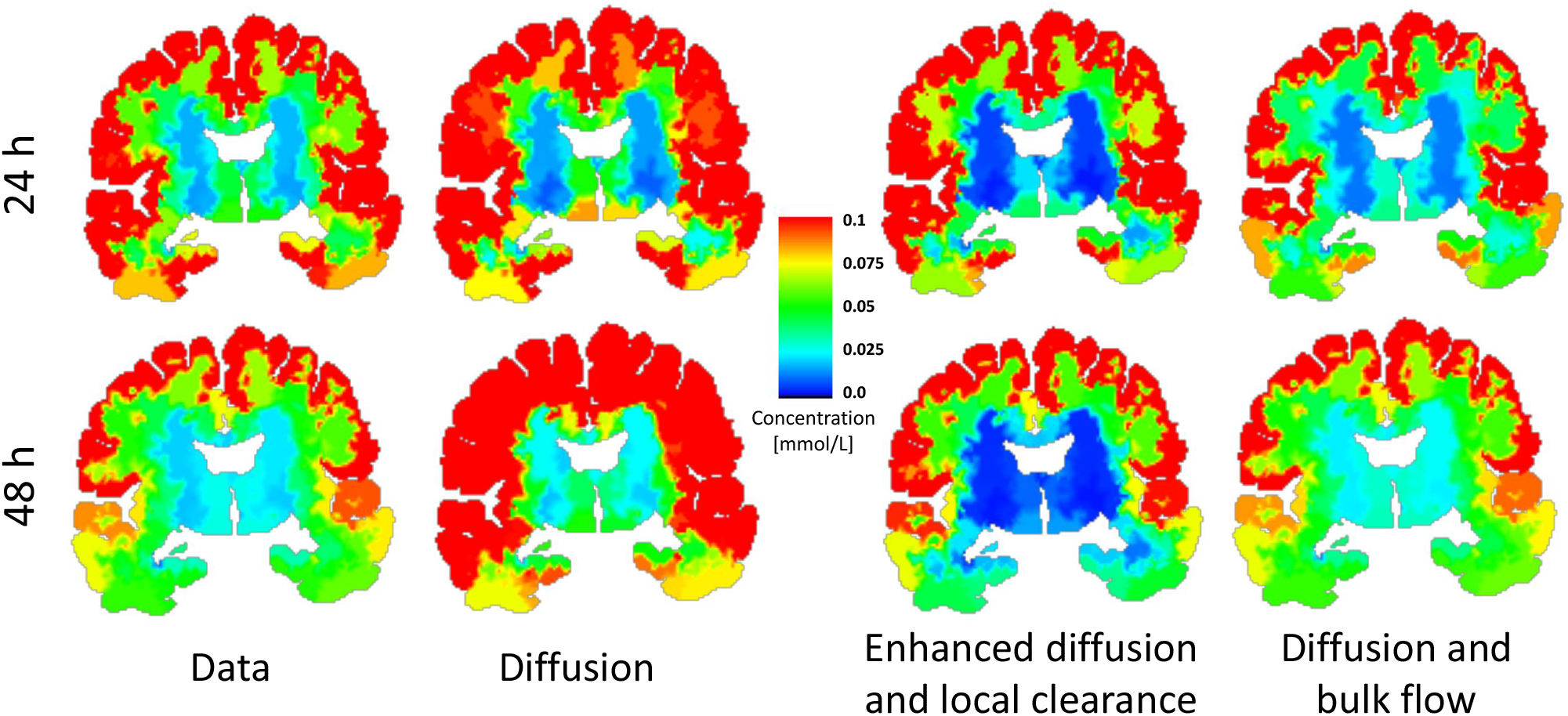
Clinical observations (data) versus simulations results for 1) pure diffusion 2) Enhanced diffusion (*α*) and local clearance (*r*) and 3) Diffusion and bulk flow (*φ*). Concentrations are shown as regional averages over all patients at 24 and 48 hours after tracer injection. All methods provide reasonable alignment with the data after 24 hours. However, 48 hours after injection, pure diffusion overestimates tracer amount found within the brain. Models using enhanced diffusion with local clearance and diffusion with bulk flow give reasonably good agreement with data both after 24 and 48 hours. Sustained concentrations in the white matter as observed in data, is not seen when local clearance is included in the model.

## Results

From multi-modal human brain MRI (T1-weighted, T1 maps, DTI) prior to intrathecal tracer injection and contrast-enhanced MRI at multiple time points 2–48 hours after^17^, we identify and quantify tracer transport characteristics using forward and inverse subject-specific computational modelling. To capture different multiscale transport mechanisms, such as diffusion in the extracellular space, dispersion due to pulsatile motion, advection by interstitial or cerebrospinal fluid flow, and flux and transport between compartments such as e.g. between the vasculature, PVS and ECS^1, 21^, we consider a diffusion-advection-reaction equation to model the tracer concentration *c = c*(*x, t*) (mmol/L) in the brain over time:

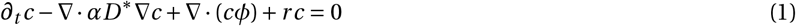

Extracellular diffusion is represented by the diffusion coefficient *D*^∗^ = *D*^∗^(*x*) of CSF tracer in brain tissue^46^, while *α* models a diffusion enhancement e.g. due to dispersion associated with pulsatile mixing (without net fluid flow)^37, 42^. The vector field *φ = φ*(*x, t*) is the velocity of an underlying fluid such as e.g. ISF bulk flow or CSF/ISF PVS flow^8, 27^, and the associated term thus models solute advection. Finally, *r* is the local clearance rate and represents solute clearance by rapid transport (molecule removal at the minute scale) e.g. via minor leakage across the blood-brain barrier (BBB)^47^ or perivascular transport.

### One-fourth of the tracers enter the brain

After intrathecal administration, tracer spreads cranially along the SAS, enters the brain tissue, and clears from the brain and SAS on a time scale of hours to days (Fig. 2A,^17^). By combining the MR signals with T_1_ maps, we quantify the tracer concentration (mmol/L) and amount of tracer (mmol) entering the brain for each subject at different time points (Fig. 2B–D). After ∼ 6 hours, nearly one-fourth (23 ± 10%, 0.116 ± 0.051 mmol) of the 0.5 mmol tracer injection has entered the brain. The maximal amount of tracers (25 ± 10%, 0.125 ± 0.050 mmol) is found within the brain after 24 hours, while 14 ± 7% (0.068 ± 0.034 mmol) remains after 48 hours. Most of the tracer enters the cerebral cortex (Fig. 2C), while less than 0.06 mmol also enters the subcortical white matter (Fig. 2D). Using subject-specific brain volumes, we find that the average concentration ranges between 0.054 - 0.377 mmol/L in the cerebral cortex and 0.014 - 0.113 mmol/L in subcortical white matter. At 24 h, the group mean is 0.146 ± 0.062 mmol/L in the cerebral cortex and 0.056 ± 0.026 mmol/L in the subcortical white matter.

**Figure 2.**
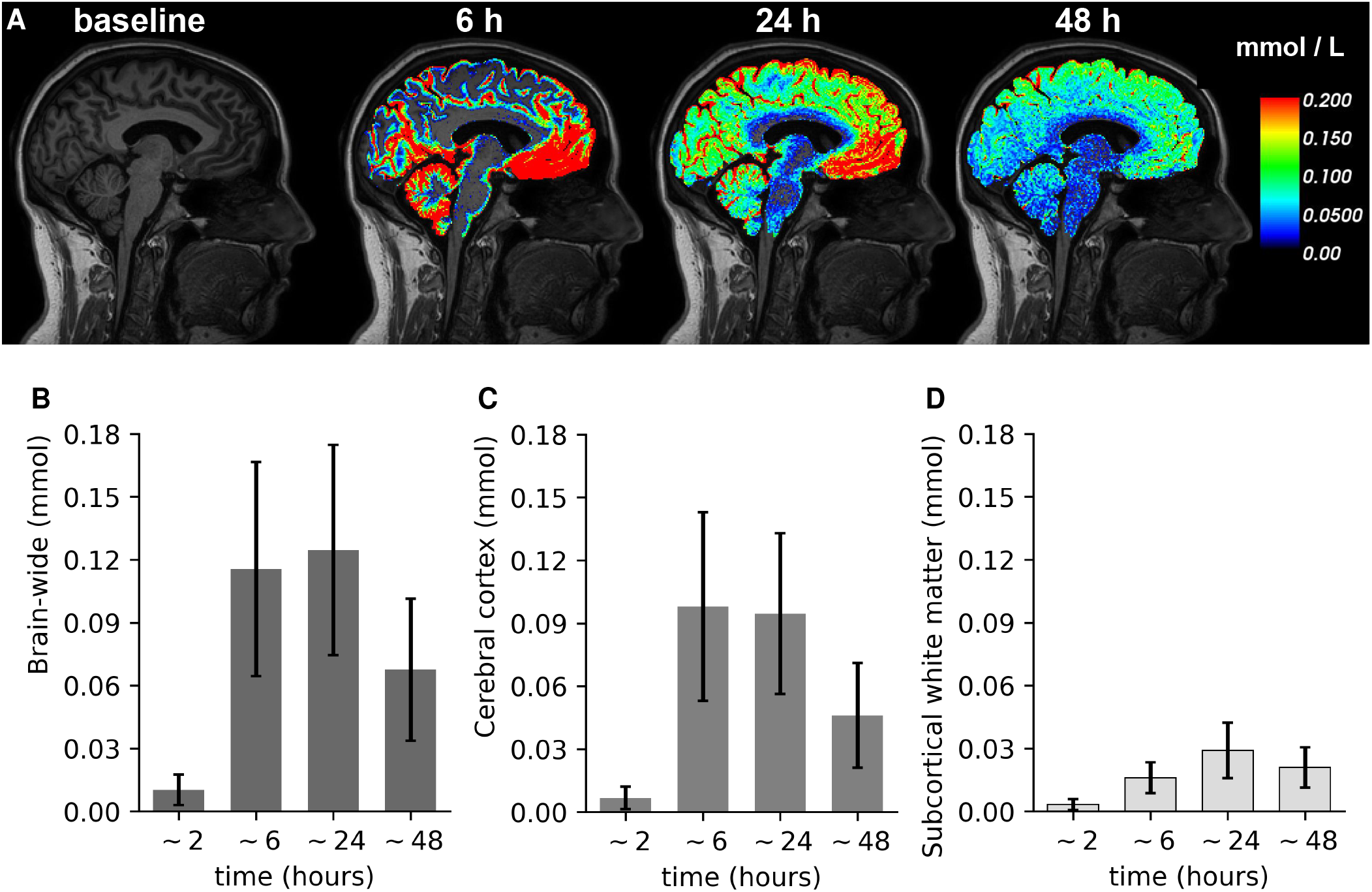
Up to 33% and on average 25% of the intrathecally injected tracer spreads into the brain over a time frame of 24–48 hours. (A) T1-weighted MRI at baseline and overlay with tracer concentrations (mmol/L) after *∼*6, 24, and 48h in a sample subject; (B)-(D): total amount of tracer (mmol) in the brain (B), cerebral cortex (C) or subcortical white matter (D) over time for all subjects combined. Error bars represent standard deviation.

### Tracer influx and clearance is more rapid than by extracellular diffusion

Accounting for baseline transport by extracellular diffusion only (*α =* 1, *r* = 0, *φ* = 0), computational predictions of tracer concentrations and amounts qualitatively agree with clinical observations (Fig. 3A,B), while key quantitative differences emerge (Fig. 3C,D). Simulated transport by extracellular diffusion underestimates the tracer influx. After ∼ 6 hours, more tracer is observed clinically than diffusion simulations predict both in the cerebral cortex (0.098 ± 0.045 vs 0.057 ± 0.029 mmol) and subcortical white matter (0.016 ± 0.007 vs 0.001 ± 0.002 mmol) (Fig. 3E, F, *α* = 1). After 24 hours, observations and simulations of tracer amounts agree in the cerebral cortex (0.095 ± 0.038 vs 0.105 ± 0.039 mmol) and subcortical white matter (0.029 ± 0.013 vs 0.038 ± 0.014 mmol). But after 48 hours, clinically observed tracer amounts were smaller compared to simulations (0.046 ± 0.025 vs 0.072 ± 0.035 mmol in the cerebral cortex and 0.021 ± 0.010 vs 0.050 ± 0.022 mmol in the subcortical white matter, Fig. 3D). Thus, extracellular diffusion alone also underestimates the tracer clearance.

**Figure 3.**
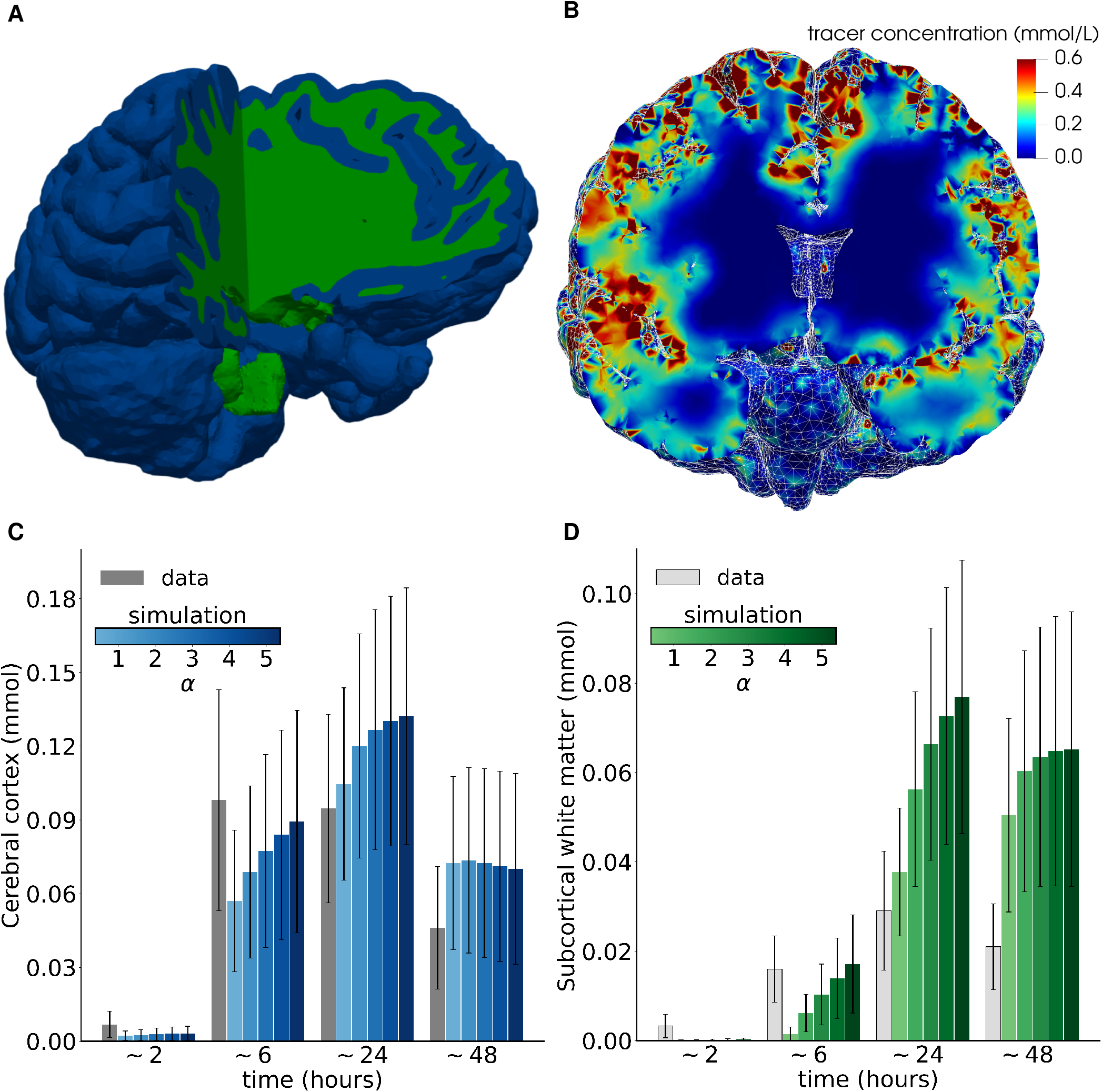
Simulations of transport by extracellular diffusion alone predict delayed influx and efflux compared to clinical observations, with delayed efflux also for enhanced diffusion. (A) Subject-specific computational brain mesh with segmentation of the cerebral cortex (blue) and subcortical white matter (green); (B) Simulated tracer distribution after 24 h in a sample subject; (C-D) Predicted versus observed average tracer concentration in the cerebral cortex (C) and subcortical white matter (D) under extracellular diffusion (*α =* 1) and enhanced diffusion (*α >* 1). Error bars represent standard deviation.

### Enhanced diffusion predicts inaccurate influx and clearance interaction patterns

Enhanced transport in pial murine PVS by cardiac-induced pulsatile mixing or CSF flow is observed in-vivo^26, 48^. Within the parenchyma, in-silico studies suggest that such enhanced transport may be modeled by an increase in the effective diffusion coefficient^28^. To investigate this effect for human brain transport, we consider effective diffusion coefficients 2–5 times greater than DTI-informed values (*α* = 2, 3, 4, 5).

As expected, the increase in diffusion coefficient accelerates tracer influx (Fig. 3C,D). In the cerebral cortex, substantially enhanced diffusion gives simulated tracer concentrations closer to (but still underestimating) clinical observations after *∼*6 hours (0.098 ± 0.045 vs 0.089 ± 0.045 mmol, *α* = 5, Fig. 3C). At 24 hours, the discrepancy increases with increasing *α*. After 48 hours, the simulated values are essentially independent of *α* in the cerebral cortex (0.07 ± 0.04 for all *α*) and around 1.5 times the tracer concentrations observed. In the subcortical white matter, similar observations hold (Fig. 3D). After 6 hours, simulations with *α* ≤ 4 underestimate the amount of tracer, while after 24 hours, simulations with *α* = 2 overestimate the data by a factor 2.1 ± 0.6 (0.029 ± 0.013 vs 0.056 ± 0.022 mmol) with increasing discrepancy for increasing *α*. Moreover, tracer concentrations in the subcortical white matter at 48 hours are overestimated by more than 2 ×. We also performed simulations with reduced diffusion coefficients yielding reduced influx, also not in agreement with the clinical observations (data not shown). Overall, these results suggest that neither diffusion nor enhanced diffusion suffice as the sole transport mechanism underlying the clinical tracer observations.

### Macroscale tracer dynamics agree with enhanced diffusion augmented by local clearance

The observation that enhanced diffusion results in more accurate estimation of tracers early on, but overestimation at later time points, suggests that a form of local clearance may occur. Turning to inverse computational modelling^34, 49^, we estimate a local clearance rate *r* > 0 and diffusion enhancement 1 < *iα* < 10 that give the best match between simulated ((1)) and observed tracer distributions for each subject. In all subjects but one, this optimization algorithm produced numerically reliable results (Tables S7, S8, S9).

The resulting models (n=23), which thus represent local clearance in addition to enhanced diffusion, agree better with the clinical data for all subjects and particularly at the 6 and 48 hour time points – both qualitatively (Fig. 1) and quantitatively in terms of the amount of tracer in the cerebral cortex and subcortical white matter (Fig. 4A,B). However, some differences in the spatial tracer distribution persist (Fig. 1). The optimal parameter configurations varied from subject to subject with *α* ∈ (1.1, 7.0) and *r* ∈ (11, 62) × 10^−4^/min, and only a weak correlation (*r =* 0.38) between *α* and *r* is found Figure 4C). The mean optimal diffusion enhancement factor was *α* = 3.5 ± 1.5, while the optimal clearance rate was *r =* (31 ± 15) × 10^−4^/min.

**Figure 4.**
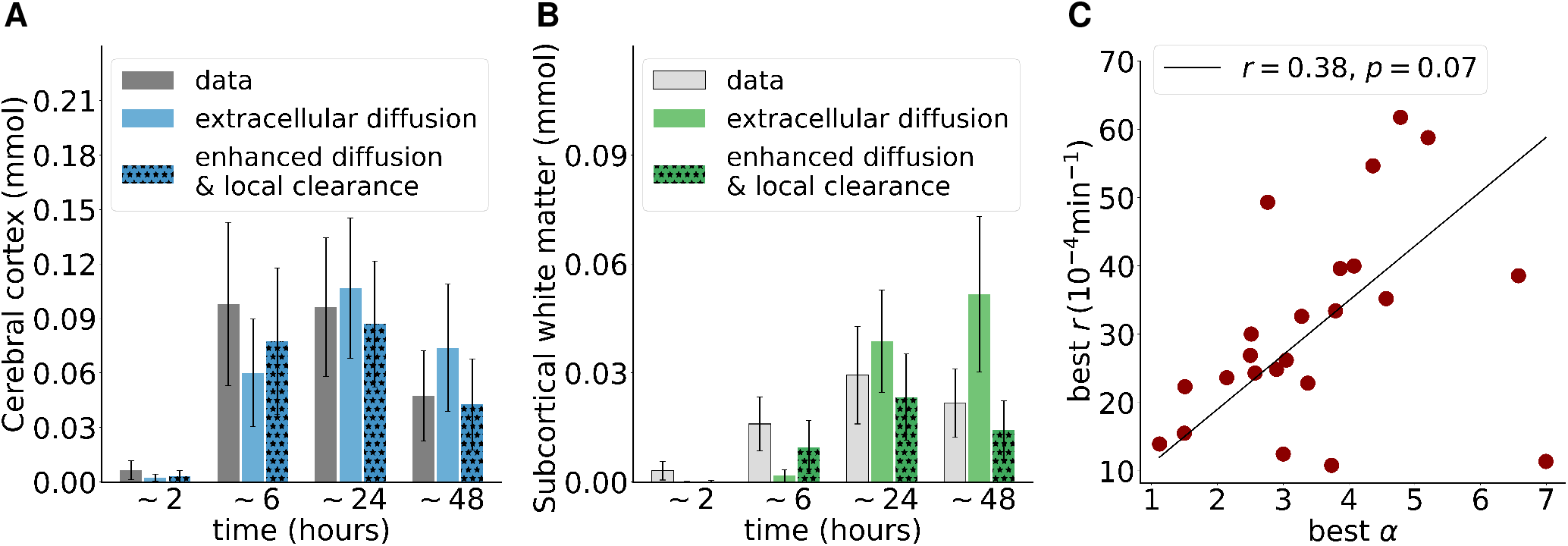
Predicted versus observed average tracer concentration in cerebral cortex (A) and subcortical white matter ((B) under diffusion and diffusion-reaction for sleep and sleep-deprived groups combined. Error bars indicate standard deviation. (C) Optimal enhanced diffusion factors *α* and local clearance rates *r*.

### Tracer patterns are compatible with extracellular diffusion augmented by advective flow at the *µ*m/min scale

The multifaceted evidence for bulk flow of interstitial or cerebrospinal fluid through the brain^27^, in particular the last decade of glymphatic research^8, 21^, naturally underpins the question of to what extent advection contributes to human brain tracer transport. Longitudinal image sets of tracer distributions allow for using high-dimensional inverse modelling to identify and quantify subject-specific flow velocity fields. More precisely, for each subject and pairs of time intervals (∼ 6 –24 and ∼ 24 – 48 hours), we estimate a spatially-varying advective velocity field *φ* that gives the minimal misfit between tracer observations and advection-diffusion simulations. Our optimization algorithm (see Methods) yielded numerically robust results for all subjects with data availability for the time interval 6–24 hours (n=22) and all but three subjects for the time interval *∼*24–48 hours (n=18, Tables S3 and S4).

The simulations reveal that the clinically observed tracer transport is compatible with persistent flow fields within the brain parenchyma at mean speeds of ∼ 1–8 *µ*m/min (Fig. 5), corresponding to average bulk flow rates of ∼ 0.02–0.16 *µ*L/(g min)^46^. The estimated velocity fields express non-trivial fluid flow patterns (Fig. 5B–C), and moderately vary between subjects, time intervals and regions. Between ∼ 6 and 24 hours, the flow speed is 2.32 ± 0.75 *µ*m/min on average brain-wide (Fig. 5D, Table S5). Flow speeds are higher in the brain stem (2.97 ± 1.30 *µ*m/min) and cerebral cortex (2.48 ± 0.81 *µ*m/min) than in the subcortical white matter (2.11 ± 0.67 *µ*m/min). Turning to the tracer clearance phase (24–48h), simulations continue to identify complex advection flow fields with brain-wide average flow speeds ranging from 1.25 to 8.39 *µ*m/min (Fig. 5E). We note that there is only a very weak (negative) correlation (∼-0.3) between the estimated average flow velocities at 6-24h and 24-48h (Fig. S6).

**Figure 5.**
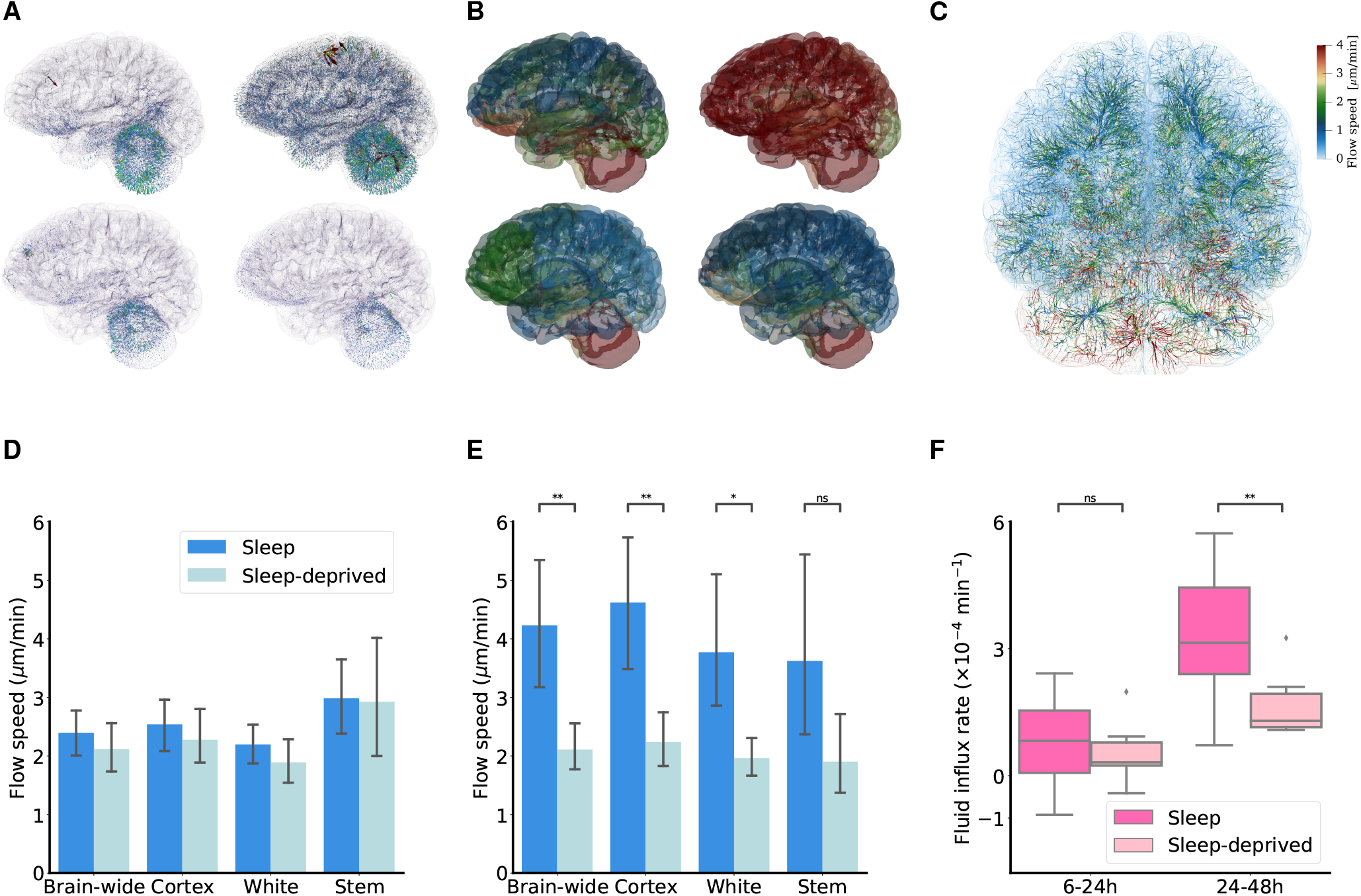
A: Advective fluid flow fields for two sample subjects (upper/lower) during 6–24h (left) and 24–48 hours (right), glyph scale: 3*×*. B: Locally (over ∼ 180 local subregions) averaged velocity field magnitudes for two sample subjects (upper/lower) during 6–24h (left) and 24–48 hours (right), scale as in C. C: Streamline visualization of sample flow field over 24–48 hours. (D) Estimated 6–24h flow speeds (velocity magnitude, see Methods) averaged brain-wide, over the cerebral cortex, over the subcortical white matter and in the brain stem for sleeping versus the sleep-deprived groups. (E) As for (D) but for 24–48h interval. (F) Brain-wide fluid influx average for the sleeping and sleep-deprived groups over 6–24h and 24-48h.

The fluid contribution to changes in tracer concentration (*∇* · (*cφ*)) is naturally decomposed into a purely advective component (*φ · ∇c*) and a volume change component (*c∇ · φ*) where the divergence of the velocity field *∇ · φ* quantifies the fluid influx rate. Computing the brain-wide average divergence of the estimated velocity fields (Fig 5F), we find that the fluid influx rate during the 6–24 hours interval is 0.72 ± 0.93 × 10^− 4^/min and higher overall in the clearance phase (24–48 hours): 2.77 ± 1.54 × 10^− 4^/min.

In summary, extracellular diffusion alone overestimates the total amount of tracer in the brain during the clearance phase (24–48 hours), enhanced extracellular diffusion combined with local clearance provides a reasonable match in the tracer distribution and amount compared to clinical data, while extracellular diffusion with DTI-based values combined with advection by an extravascular flow field varying in space and time gives a near perfect match (Fig. 1). We do however note that of the two inverse modelling approaches, the former uses only two control variables, while the advective field allows for much more variation.

### Tracer clearance by advection is reduced in sleep deprivation group

Seven of the 24 subjects underwent total sleep deprivation during the first 24 hours after injection. The MRI signal intensity of these subjects differ compared to sleeping controls, as previously reported^17^. To quantify, we here find that during the early influx phase (0–6 hours), the amount of tracer within the brain is comparable for the two groups (0.111 ± 0.056 mmol vs 0.127 ± 0.030 mmol, Student’s t-test, p=0.54). However, higher amounts of tracer linger in the brain of the sleep deprived subjects after 48 hours (0.057 ± 0.030 mmol vs 0.088 ± 0.031 mmol, Student’s t-test, p=0.0495), with the larger concentration differences in the subcortical white matter (0.018±0.009 mmol vs 0.028±0.006 mmol, Student’s t-test, p=0.027). Importantly, to assess whether differences between the groups stem from differences in tracer availability at the pial surface, we also evaluated the total amount of tracer (per unit depth) over the brain surface for each group: these differences were found to be non-significant (p=0.55, 0.40, 0.14 for ∼ 6, 24, 48 hours after injection, respectively).

One can then ask whether the effect of sleep deprivation is also captured by the tracer transport parameters; i.e. enhanced diffusivity, local clearance, or advective flow velocity? Interestingly, the delayed tracer clearance after sleep-deprivation is captured by marked differences between the groups in terms of fluid flow velocities and thus advection over the 24–48h time interval (Fig. 5E). Flow speeds are lower by nearly a factor two in the sleep-deprived (*n* = 7) compared to the sleep (*n* = 11) group (2.11 ± 0.59 *µ*m/min vs 4.23 ± 1.98 *µ*m/min, Welch’s t-test, p=0.0057) brain-wide, in the cerebral cortex (2.24 ± 0.69 *µ*m/min vs 4.62 ± 2.03 *µ*m/min, p=0.0033), and in the subcortical white matter (1.96 ± 0.47 *µ*m/min vs 3.77 ± 1.98 *µ*m/min, p=0.014). Moreover, the reduced flow speeds are mirrored by reduced fluid influx rates between 24 and 48 hours: 1.68 ± 0.79 ×10^−4^/min for the sleep-deprived vs 3.48 ± 1.51 ×10^−4^/min (Welch’s t-test, p=0.0051) for the sleep group. On the other hand, differences in enhanced diffusion (*α*) and clearance (*r*) between the sleep (*α* = 3.7 ± 1.6, *r =* (34 ± 14) × 10^−4^/min) and sleep deprived group (*α* = 2.9 ± 0.9, *r* = (23 ± 11) × 10^−4^/min) were found to be non-significant (*p* = 0.22 and *p* = 0.11 for *α* and *r*, respectively).

## Discussion

Our findings support the notion that extracellular diffusion alone is not sufficient as a brain-wide tracer transport mechanism. Instead, we show that human MRI observations align well with transport by either (i) substantially enhanced (3.5*×*) extracellular diffusion in combination with local clearance rates corresponding to a tracer half-life of up to 5 hours, (ii) or extracellular diffusion augmented by advection with advective ISF or CSF/ISF flow speeds of 1–8 *µ*m/min on average. The estimated local clearance rates and the flow speeds are within the range reported in the literature (as detailed below), while the effective enhancement is much larger than what is previously reported in humans while still lower than that reported in mice.

Our quantification of tracer transport into the brain parenchyma reveals that 23–25% ± 10% of the injected tracers were found within the brain between 6 and 24 hours. The peak tracer concentrations in the cerebral cortex and subcortical white matter compare well with the respective concentrations reported by Watts et al in a single subject^30^. These results, in conjunction with tracer concentrations in the cranial CSF^30^, suggest that no more than 33% of injected tracers are located within the intracranial compartment at any given point in time; conversely, that 67% of the injected tracer has not passed from the intrathecal to the intracranial compartment. It is not possible to quantify spinal versus cranial outflow from these numbers, however they suggest that a dominant unidirectional flow directed towards the upper convexities of the brain, as suggested by Cushing’s third circulation^50^, is unlikely. Previous studies have suggested that 15–35% of CSF is drained along the spinal cord^51–53^, mainly via spinal nerve roots^54^. The present data suggest that up to 70% is drained directly from the thecal sac. These observations also highlight the importance of accounting for flow and transport within the CSF compartment, or via the availability of tracer at the pial surface as a proxy, when quantifying brain tracer influx^16^.

Diffusive transport within the interstitium is expected to dominate transport over short distances^5, 23, 24, 44^. However, at the scale of the human brain, previous studies support the presence of additional transport mechanisms such as enhanced diffusion^34^ or directional flow^33^, although the additional transport needed is relatively small. In mice on the other hand, Ray et al.^28^ found that diffusive transport with effective diffusion coefficients 10–25 times greater than extracellular diffusion could explain parenchymal tracer transport. It should be noted that the dynamics of gadolinium-based contrast agent transport are much faster in rodents where peak concentration within the brain occurs around one hour after injection into the cisterna magna^55^. In addition, mice have a CSF turnover time three times shorter than humans^56^, suggesting higher fluid velocities in the SAS. Recently, we have also shown that higher SAS velocities reduce time to peak concentration in the parenchyma^57^. Different SAS dynamics may thus at least partially explain the discrepancy between estimated dispersion coefficients in mice^28^ versus humans, both in previous^34^ and present studies. Interestingly, as shown here enhanced diffusion may explain clinical observations during the influx phase (t < 24 hours), but not during the outflux phase.

A scenario with local clearance with a clearance rate *r* combined with enhanced diffusion gives reasonable match with clinical observations, not only brain-wide but also in the cerebral cortex and in subcortical white matter. The cohort average *α* = 3.5 corresponds to an effective diffusion coefficient of *D*eff = 3.25 × 10^−4^ mm^2^ s^−1^. This value is close to the free diffusion coefficient of gadobutrol in water *D* = 3.8 *×* 10^*−*4^ mm^2^s^−134^. Interpreting the results as enhanced diffusion thus suggest a tortuosity of 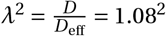. In the literature, based on experimental diffusion data *λ* ∼ 1.6^46^, though the geometrical (*λ* = 1.18) and viscous (*λ =* 1.20) components of the tortuosity are much smaller^22^ and not sufficient to account for *λ* = 1.6. Moreover, the estimated local clearance rate of *r* = 31 ± 15 × 10^−4^/min corresponds to an exponential decay half-life of log 2/*r* = 287 minutes. The clearance pathways described by this parameter relate to minor leakage across the blood-brain-barrier, or very rapid transport along paravascular pathways at scales not detectable with MRI. As such, our clearance rate estimate could be compared to the half-life of gadobutrol to the blood (from the subarachnoid space) of 3.83 hours or 230 minutes as reported by Hovd et al.^47^. Also note that the subject variation is similar when comparing the clinical data (ibid) and our estimation (150 versus 153 min). The clearance is comparable although somewhat larger than blood-brain leakage observed in dementia^58^ of 1 − 20 × 10^−4^/min.

The 2004-review by Abbott^27^, in part based on experimental research by Cserr, Rosenberg and coauthors during the 1980’s^59–62^, highlights “clear evidence for the presence of a bulk flow of brain ISF at a rate of 0.1–0.3 *µ*L/(g min)”. Nicholson^4^ interprets the same experimental studies to support bulk flow velocities of 5.5–14.5 *µ*m/min. Our values are in remarkable agreement with these classical estimates: modelling human tracer movement as governed by extracellular diffusion in combination with advection by fluid flow with average flow speeds of 1.25–8.39 *µ*m/min gives excellent agreement with clinical data. The estimated flow speeds are higher in regions around the cerebellum, in agreement with relative results reported by Koundal et al.^44^. As the estimated flow speeds represent volume averages, we may use porous media theory to estimate corresponding average local velocities. First, assuming that PVSs occupy 1% of the brain volume, and ignoring interstitial velocities, our findings are compatible with average PVS velocities of 2.1–14.0 *µ*m/s, i.e. velocities of the same order or somewhat lower than in pial PVS in mice^26^. On the other hand, ignoring PVS flow and assuming an extra-cellular volume fraction of 15%, we obtain average extracellular velocities of 0.14–0.92 *µ*m/s, which are in line with the analysis of Ray et al.^63^, but 1–2 orders higher than the upper estimates reported by Holter et al.^24^.

The advective flow fields identified via high-dimensional inverse modelling admit, and are indeed supported by, a non-trivial and non-vanishing average fluid influx rate on the order of −0.5–4 × 10^−4^/min. Interestingly, this net fluid production changes from the tracer influx phase (low fluid influx or even fluid outflux) to the tracer clearance phase (higher fluid influx) suggesting that flow always contributes to speed up the transport. This is not in line with a constant production/filtration over capillaries^64^, where a positive fluid influx would be expected. Comparing our inverse flow estimation approach with the optimal mass transport methods introduced and refined by Tannenbaum, Benveniste and coauthors^43–45^, we emphasize that we here explicitly include extracellular diffusion in the underlying transport problem, directly use the velocity field to represent advection rather than e.g. anisotropic diffusion, allow for local fluid influx/efflux, target numerical robustness by simultaneous approximation of the velocity and concentration fields, and provide quantitative (absolute) flow field estimates.

CSF and glymphatic function change according to circadian rhythm and/or sleep in animal models. In particular, extra-cellular volume fraction, perivascular intake and interstitial clearance^15^, lymphatic efflux^16^, choroid plexus gene expression^65^, AQP4 polarization and drainage to lymph nodes^66^, perivascular pulsations^67^ all display significant variations. In humans, much less is known about CSF flow and exchange. However,^68^ demonstrated a direct link between CSF dynamics, hemodynamics, and neural activity during sleep. Further, in^17^, CSF tracer distribution differed in subjects that were sleeping and sleep-deprived, most notably after 48 hours and in particular in subregions such as the limbic system. Based on our modeling, we find that the average advective velocity here is nearly halved in sleep-deprived and the differences between the groups are statistically significant. For the enhanced diffusion and clearance parameters we found differences but they were not statistically significant.

From a mathematical point of view, the forward diffusion-advection-reaction problem (1) is a well-posed problem. This implies that solutions are unique, and stable in the sense that small variations in input data give only small variations in the output quantities. Moreover, the numerical methods used are guaranteed to provide numerically reliable approximations. In contrast, for the inverse optimization problems, there may be multiple solutions (*r, α*) or (*φ, c*) that give an optimal fit to the MRI data and satisfy (1), and minor variations in the data (e.g. due to noise) may strongly affect the estimated parameters. For instance, the (*α, r*) parameters may cancel out in the sense that a high *α* combined with a high *r* may (in certain regions depending on the scales in space and time and in an average sense) yield similar transport dynamics as combinations of smaller *α, r*. Moreover, we cannot guarantee that a global optimum is attained, but rather that the parameters are optimal in a local neighborhood of potential values. The high-dimensional velocity field estimation may also be prone to over-fitting with regard to noisy and relatively sparse data.

These challenges are addressed by the presence and choice of regularization terms in the objective functional (2), and to some extent the box constraints associated with *r* and *α*. These are well-established and well-posed approaches to regain uniqueness and stability for the inverse models^49, 69^. Furthermore and importantly, the fact that our computational technology yields numerically reliable solutions (that are robust with respect to numerical parameter choices such as time steps and regularization parameters) for the vast majority of simulation cases is promising. We also note that the correlation between *α* and *r* is weak (correlation coefficient 0.38). We also remark while one may always seek more advanced modelling, computing the results reported in this study required around 64,000 CPU hours and can as such be thought of as a reasonable and feasible compromise in terms of complexity.

In addition to the inverse modelling aspects discussed above, we note that boundary conditions were prescribed by a linear interpolation between data points in time, and projected directly onto the finite element mesh. In data from Watts et al.^30^, the peak concentration occurs at around 10 hours in the CSF and at closer to 15 hours in the cerebral cortex. Even though the report from Watts^30^ only considered a single subject, there is a risk that we miss the point of peak concentration in our data set. However, an extra measurement with peak in CSF concentration at ∼ 10 hours would not alter our observation that extracellular diffusion is too slow to explain the measurements at ∼ 6 hours, and would only increase the amount of tracer at ∼ 48 hours predicted by a pure extracellular diffusion model. We also note that our flow field estimation cannot resolve non-linear velocity variations over distances shorter than the computational mesh size (a few mm), and that velocities may be underestimated in regions where there is little or no tracer present at any time point.

In the midst of a wave of neuroimaging advances across scales^21, 70^, here high-fidelity inverse computational models create a bridge between multi-modal MR imaging data and biophysical clearance hypotheses; thus enabling a new technological avenue for identification and quantification of human brain solute transport mechanisms. Our findings highlight the combined roles and importance of extracellular diffusion, local clearance at rates comparable to tracer transport across the blood-brain barrier or advective velocities on the order of *µ*m/min sustained by local fluid influx or efflux, and reveal reduced advective flow after sleep-deprivation. Distinguishing between these clearance mechanisms calls for new clinical or experimental protocols combining in-vivo brain imaging with blood, lymph and crucially CSF measurements.

## Methods

### Data collection and approvals

In reference (sleep) (*n* = 17) and sleep-deprivation (*n* = 7) subject groups, *T*_1_-weighted MRI, *T*_1_-maps and DTI were collected prior to intrathecal injection of CSF tracer (gadobutrol), while contrast-enhanced MR images were collected at multiple time points between 0 and 48 hours post injection, as previously reported_17_. During the night between day 1 and 2 (12–24h post injection), individuals in the sleep-deprived group were deprived of sleep, while the reference group slept as normal. The study_17_ was approved by the Regional Committee for Medical and Health Research Ethics (REK) of Health Region South-East, Norway (2015/96), the Institutional Review Board of Oslo University Hospital (2015/1868), the National Medicines Agency (15/04932-7), and was registered in Oslo University Hospital Research Registry (ePhorte 2015/1868). The conduct of the study was governed by ethical standards according to the Declaration of Helsinki of 1975 (and as revised in 1983). Study participants were included after written and oral informed consent.

### Computational geometries

For each subject, we generate subject-specific 3D meshes at different resolution levels (low-res, standard, high-res) using FreeSurfer_71_ and SVMTK_72_. A typical standard (high-res) mesh Ω consists of 1.1 (4.2) million tetrahedral mesh cells of diameter 0.4 − 5.5 (0.1 − 2.8) mm (Table S1). We define and label the cerebral cortex, subcortical white matter and brain stem as disjoint regions within the mesh via the pial surface, white-gray matter interface, and subdomain tags generated by FreeSurfer (Fig. 3A). In addition to these larger regions, we identified and labeled nearly 180 smaller brain subregions using the FreeSurfer parcellation.

### Mapping signal intensities to concentrations

Contrast-enhanced signal intensities may be mapped to tracer concentrations via a map of the (spatially-varying) relaxation times *T*_1_, as previously described_34_. For 15 of the 24 subjects, subject-specific *T*_1_ maps were measured during data collection, while for the remaining subjects, such were not available. To compensate while avoiding introducing bias between groups, we used group-averaged and regionally constant *T*_1_ values for all subjects. This method was compared against using raw *T*_1_ maps or filtered *T*_1_ maps for the 15 subjects with *T*_1_ maps available, with the different approaches yielding tracer concentration values that differed by at most 11% (Section S1.1).

### Tracer transport equations

We model the concentration *c*(*x, t*) (in mmol/L) of CSF tracer as a function of time *t* > 0 and space *x* ∈ Ω solving (1), and thus distributing via three modes of transport: diffusion with a heterogeneous diffusion coefficient *D*^∗^ = *D*^∗^(*x*) and enhancement factor *α* > 1, advection via a velocity vector field *φ* = *φ*(*x, t*) ∈ **R**_3_ and local clearance at a clearance rate *r* > 0. On the boundary and for each time *t* > 0, we prescribe the observed CSF tracer concentrations *c*_mri_(*x, t*), mapped from the MRI signal intensities as described above and linearly interpolated in time between MRI scans. As initial condition at *t* = *t*_0_, corresponding to the baseline image, we set CSF tracer concentration in the entire domain to be *c*(*·, t*_0_) = 0. For all time points *t*, we compute the amount of tracer *M*_*i*_ (*t*) and the average concentration 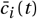) in all FreeSurfer-labeled regions as well as the average concentration in the cerebral cortex, subcortical white matter, and brain-wide by integrating over the respective regions; that is, for each region Ω_*i*_:

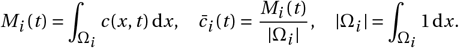

The amount of tracer per unit area on the brain surface *∂*Ω was computed for all subjects as

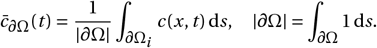

### Diffusion and dispersion

Tracer transport by diffusion only is represented by letting *α* = 1, *φ* = 0, *r* = 0 in (1). The anisotropic and spatially-varying diffusion tensor *D*^∗^ can be estimated from DTI. However for 6 of the 24 subjects, no DTI data were available. For the 18 subjects with DTI, we solved (1) with (a) subject-specific *D* estimated voxel-wise from DTI as well as (b) group-averaged and regionally varying isotropic (scalar) diffusion parameters. The two methods differed by about 1%. We therefore used the latter method for all 24 subjects and all forward and inverse simulations (Section S1.2).

Enhanced diffusion, for instance via dispersion_28, 34, 73_, is represented via the enhancement factor *α* > 1. Specifically, we consider enhanced diffusion-scenarios for which *α* = 1, 2, 3, 4, 5 in the cerebral cortex while keeping *α* = 1 fixed in the subcortical white matter.

### Numerical methods and software

The CSF tracer concentrations were represented as continuous piecewise linear polynomials defined over the computational mesh(es) via interpolation. For each subject and each set of model parameter variations, we solve the diffusion-advection-reaction equation (1) from *t*_0_ to *T* (or for a single time window between consecutive MR scans at *t*_1_ and *t*_2_) using a second-order finite difference scheme in time and a finite element method yielding second-order approximations in space of the concentration field via the FEniCS finite element software_74_. The standard resolution meshes yield results that differ at most 4% to the high resolution meshes for the forward simulations (Section S2.2) and were therefore used for the reported results. The simulation end time *T* (∼ 48 hours) was set as the time of the last MR scan for each subject. All computations were performed on resources provided by Sigma2 - the National Infrastructure for High Performance Computing and Data Storage in Norway.

### Inverse identification of an advective velocity field

To identify an underlying velocity field *φ* that match the tracer observations as well as the biophysics described by (1), we adapt and apply an inverse problem technique_69_. For any pair of MRI scans (at *t*_1_ and *t*_2_ with *τ = t*_2_ − *t*_1_ (hours)), we map the CSF tracer observations *c*_mri_(*t*_1_) and *c*_mri_(*t*_2_) onto the computational mesh. For each subject and each such time interval [*t*_1_, *t*_2_], we then consider the following constrained optimization problem: find a spatially-varying velocity field *φ* (*φ*(*x*) ∈ **R**_3_, *x* ∈ Ω) that minimizes the discrepancy between simulated and observed concentrations and is sufficiently

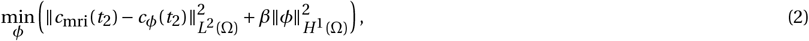

and is such that *c* = *c*_*φ*_ numerically solves (1) with this *φ* from *t*_1_ to *t*_2_ with timestep *τ*, with the observations *c*_mri_(*t*_1_) prescribed as the initial condition at *t*_1_, and with *c*_mri_(*t*_2_) prescribed as boundary conditions at *t*_2_. Here, *I· I*_*L*_2 denote the standard *L*_2_(Ω)-norm, and similarly for *H* _1_(Ω)_75_, while *β* = 10_−4_ is a regularization parameter enforcing the additional smoothness of the solution *φ*. This *φ* is thus a quantification of an underlying fluid flow field that may transport the tracer by advection such as e.g. ISF flow or an averaged representation of a more localized (PVS) flow.

We solve this high-dimensional optimization problem using a reduced approach with a maximum of 80 iterations of the L-BFGS optimization algorithm_76_ as implemented in the Dolfin-adjoint software_77_ using FEniCS_74_ and SciPy_78_. Specifically, we compute velocity field predictions for during the first day (∼ 1 –6h), the first evening/night (∼ 6–24h) and day 2 (∼ 24–48h). The estimated velocity fields were stable with respect to variations in mesh resolution and regularization parameters (Section S3.1).

### Flow and velocity quantities of interest

To quantify the fluid flow, we compute average flow speeds as the velocity magnitude field averaged brain-wide:

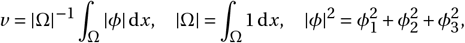

or instead averaged over larger regions (cerebral cortex, subcortical white matter and brain stem) or smaller regions (defined by subject-specific FreeSurfer parcellations). The local fluid influx or efflux described by any velocity field *φ* = (*φ*_1_, *φ*_2_, *φ*_3_) is given by its divergence:

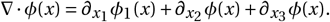

### Local clearance rates

Assuming that local clearance of molecules, e.g., via the microcirculation or cellular degradation, is proportional to their concentration *c* yields the clearance/reaction termr *c* in (1).

In order to determine subject-specific optimal dispersion and reaction constants (*α, r*) we formulate the following PDE constrained optimization

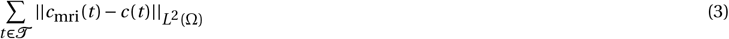

where *c*_*mri*_(*t*) is the tracer concentration estimated from MRI at times *t* ∈ 𝒯 and *c*(*t*) is the numerical solution to (1) with *ϕ* = 0. The initial and boundary conditions are the same as in (1) and we fix *α* = 1 in the white matter We constrain *α* ∈ [1,10] and *r* ∈ [10^−7^ s^−1^,10^−3^ s^−1^]. This problem is then solved for time step sizes of 30, 20, 10 and 5 min (Table S9), and the combination of parameters (*α, r*) found by the minimization algorithm with the smallest time step tested (either 10 or 5 min) reported.

## Acknowledgements

M. E. Rognes acknowledges support and funding from the European Research Council (ERC) under the European Union’s Horizon 2020 research and innovation programme under grant agreement 714892, and the Research Council of Norway (RCN) via FRIPRO grant agreement #324239 (EMIx). K.-A. Mardal acknowledges support from the Research Council of Norway, Grant 300305 and 301013, and the national infrastructure for computational science in Norway, Sigma2, Grant NN9279K.

## Author contributions statement

Study design: VV, BZ, GR, PKE, MER, KAM. Data collection: GR, PKE. Method development: VV, BZ, MER, KAM. Simulations: VV, BZ, MER, KAM. All authors contributed to the drafting and editing of the manuscript.

## References

1. Tarasoff-Conway, J. M. et al. Clearance systems in the brain—implications for alzheimer disease. Nat. reviews neurology 11, 457–470 (2015).

2. Matsumae, M. et al. Research into the physiology of cerebrospinal fluid reaches a new horizon: intimate exchange between cerebrospinal fluid and interstitial fluid may contribute to maintenance of homeostasis in the central nervous system. Neurol. medico-chirurgica 56, 416–441 (2016).

3. Louveau, A. et al. Understanding the functions and relationships of the glymphatic system and meningeal lymphatics. The J. clinical investigation 127, 3210–3219 (2017).

4. Nicholson, C. Diffusion and related transport mechanisms in brain tissue. Reports on progress Phys. 64, 815 (2001).

5. Smith, A. J. & Verkman, A. S. Going against the flow: Interstitial solute transport in brain is diffusive and aquaporin-4 independent. The J. physiology 597, 4421 (2019).

6. Cserr, H. F. & Ostrach, L. Bulk flow of interstitial fluid after intracranial injection of blue dextran 2000. Exp. neurology 45, 50–60 (1974).

7. Rennels, M. L., Gregory, T. F., Blaumanis, O. R., Fujimoto, K. & Grady, P. A. Evidence for a ‘paravascular’fluid circulation in the mammalian central nervous system, provided by the rapid distribution of tracer protein throughout the brain from the subarachnoid space. Brain research 326, 47–63 (1985).

8. Iliff, J. J. et al. A paravascular pathway facilitates csf flow through the brain parenchyma and the clearance of interstitial solutes, including amyloid β. Sci. translational medicine 4, 147ra111–147ra111 (2012).

9. Harrison, I. F. et al. Impaired glymphatic function and clearance of tau in an alzheimer’s disease model. Brain 143, 2576–2593 (2020).

10. Ringstad, G. et al. Brain-wide glymphatic enhancement and clearance in humans assessed with mri. JCI insight 3 (2018).

11. Gaberel, T. et al. Impaired glymphatic perfusion after strokes revealed by contrast-enhanced mri: a new target for fibrinolysis? Stroke 45, 3092–3096 (2014).

12. Thrane, A. S., Thrane, V. R. & Nedergaard, M. Drowning stars: reassessing the role of astrocytes in brain edema. Trends neurosciences 37, 620–628 (2014).

13. Mehta, A. I., Linninger, A., Lesniak, M. S. & Engelhard, H. H. Current status of intratumoral therapy for glioblastoma. J. neuro-oncology 125, 1–7 (2015).

14. Lohela, T. J., Lilius, T. O. & Nedergaard, M. The glymphatic system: implications for drugs for central nervous system diseases. Nat. Rev. Drug Discov. 21, 763–779 (2022).

15. Xie, L. et al. Sleep drives metabolite clearance from the adult brain. science 342, 373–377 (2013).

16. Ma, Q. et al. Rapid lymphatic efflux limits cerebrospinal fluid flow to the brain. Acta neuropathologica 137, 151–165 (2019).

17. Eide, P. K., Vinje, V., Pripp, A. H., Mardal, K.-A. & Ringstad, G. Sleep deprivation impairs molecular clearance from the human brain. Brain 144, 863–874 (2021).

18. Wardlaw, J. M. et al. Perivascular spaces in the brain: anatomy, physiology and pathology. Nat. Rev. Neurol. 16, 137–153 (2020).

19. Hladky, S. B. & Barrand, M. A. The glymphatic hypothesis: the theory and the evidence. Fluids Barriers CNS 19, 1–33 (2022).

20. Rasmussen, M. K., Mestre, H. & Nedergaard, M. Fluid transport in the brain. Physiol. Rev. 102, 1025–1151 (2022).

21. Kelley, D. H. et al. The glymphatic system: Current understanding and modeling. Iscience 104987 (2022).

22. Nicholson, C. & Hrabětová, S. Brain extracellular space: the final frontier of neuroscience. Biophys. journal 113, 2133–2142 (2017).

23. Jin, B.-J., Smith, A. J. & Verkman, A. S. Spatial model of convective solute transport in brain extracellular space does not support a “glymphatic” mechanism. J. Gen. Physiol. 148, 489–501 (2016).

24. Holter, K. E. et al. Interstitial solute transport in 3D reconstructed neuropil occurs by diffusion rather than bulk flow. Proc. Natl. Acad. Sci. 201706942 (2017).

25. Wentland, A. L., Wieben, O., Korosec, F. R. & Haughton, V. M. Accuracy and reproducibility of phase-contrast mr imaging measurements for csf flow. Am. J. Neuroradiol. 31, 1331–1336 (2010).

26. Mestre, H. et al. Flow of cerebrospinal fluid is driven by arterial pulsations and is reduced in hypertension. Nat. communications 9, 1–9 (2018).

27. Abbott, N. J. Evidence for bulk flow of brain interstitial fluid: significance for physiology and pathology. Neurochem. international 45, 545–552 (2004).

28. Ray, L. A., Pike, M., Simon, M., Iliff, J. J. & Heys, J. J. Quantitative analysis of macroscopic solute transport in the murine brain. Fluids Barriers CNS 18, 1–19 (2021).

29. Ringstad, G. et al. Non-invasive assessment of pulsatile intracranial pressure with phase-contrast magnetic resonance imaging. PloS one 12, e0188896 (2017).

30. Watts, R., Steinklein, J., Waldman, L., Zhou, X. & Filippi, C. Measuring glymphatic flow in man using quantitative contrast-enhanced mri. Am. J. Neuroradiol. 40, 648–651 (2019).

31. Eide, P. K., Valnes, L. M., Lindstrøm, E. K., Mardal, K.-A. & Ringstad, G. Direction and magnitude of cerebrospinal fluid flow vary substantially across central nervous system diseases. Fluids Barriers CNS 18, 1–18 (2021).

32. Eide, P. K. et al. Altered glymphatic enhancement of cerebrospinal fluid tracer in individuals with chronic poor sleep quality. J. Cereb. Blood Flow & Metab. 0271678X221090747 (2022).

33. Croci, M., Vinje, V. & Rognes, M. E. Uncertainty quantification of parenchymal tracer distribution using random diffusion and convective velocity fields. Fluids Barriers CNS 16, 1–21 (2019).

34. Valnes, L. M. et al. Apparent diffusion coefficient estimates based on 24 hours tracer movement support glymphatic transport in human cerebral cortex. Sci. reports 10, 1–12 (2020).

35. Kinney, J. P. et al. Extracellular sheets and tunnels modulate glutamate diffusion in hippocampal neuropil. J. Comp. Neurol. 521, 448–464 (2013).

36. Daversin-Catty, C., Vinje, V., Mardal, K.-A. & Rognes, M. E. The mechanisms behind perivascular fluid flow. bioRxiv (2020).

37. Sharp, M. K., Carare, R. O. & Martin, B. A. Dispersion in porous media in oscillatory flow between flat plates: applications to intrathecal, periarterial and paraarterial solute transport in the central nervous system. Fluids Barriers CNS 16, 13 (2019).

38. Kedarasetti, R. T., Drew, P. J. & Costanzo, F. Arterial pulsations drive oscillatory flow of csf but not directional pumping. Sci. reports 10, 1–12 (2020).

39. Kedarasetti, R. T. et al. Functional hyperemia drives fluid exchange in the paravascular space. Fluids Barriers CNS 17, 1–25 (2020).

40. Thomas, J. H. Fluid dynamics of cerebrospinal fluid flow in perivascular spaces. J. Royal Soc. Interface 16, 20190572 (2019).

41. Tithof, J., Kelley, D. H., Mestre, H., Nedergaard, M. & Thomas, J. H. Hydraulic resistance of periarterial spaces in the brain. Fluids Barriers CNS 16, 19 (2019).

42. Troyetsky, D. E., Tithof, J., Thomas, J. H. & Kelley, D. H. Dispersion as a waste-clearance mechanism in flow through penetrating perivascular spaces in the brain. Sci. reports 11, 1–12 (2021).

43. Ratner, V. et al. Cerebrospinal and interstitial fluid transport via the glymphatic pathway modeled by optimal mass transport. Neuroimage 152, 530–537 (2017).

44. Koundal, S. et al. Optimal mass transport with lagrangian workflow reveals advective and diffusion driven solute transport in the glymphatic system. Sci. reports 10, 1–18 (2020).

45. Benveniste, H. et al. Glymphatic cerebrospinal fluid and solute transport quantified by mri and pet imaging. Neuroscience 474, 63–79 (2021).

46. Nicholson, C. Diffusion and related transport mechanisms in brain tissue. Reports on progress Phys. 64, 815 (2001).

47. Hovd, M. H. et al. Population pharmacokinetic modeling of csf to blood clearance: prospective tracer study of 161 patients under work-up for csf disorders. Fluids Barriers CNS 19, 1–14 (2022).

48. Bedussi, B., Almasian, M., de Vos, J., VanBavel, E. & Bakker, E. N. Paravascular spaces at the brain surface: Low resistance pathways for cerebrospinal fluid flow. J. Cereb. Blood Flow & Metab. 38, 719–726 (2018).

49. Hinze, M., Pinnau, R., Ulbrich, M. & Ulbrich, S. Optimization with PDE constraints, vol. 23 (Springer Science & Business Media, 2008).

50. Cushing, H. et al. The third circulation and its channels. Lancet 2, 851–857 (1925).

51. Edsbagge, M., Tisell, M., Jacobsson, L. & Wikkelso, C. Spinal csf absorption in healthy individuals. Am. J. Physiol. Integr. Comp. Physiol. 287, R1450–R1455 (2004).

52. Marmarou, A., Shulman, K. & Lamorgese, J. Compartmental analysis of compliance and outflow resistance of the cerebrospinal fluid system. J. neurosurgery 43, 523–534 (1975).

53. Bozanovic-Sosic, R., Mollanji, R. & Johnston, M. Spinal and cranial contributions to total cerebrospinal fluid transport. Am. J. Physiol. Integr. Comp. Physiol. 281, R909–R916 (2001).

54. Proulx, S. T. Cerebrospinal fluid outflow: a review of the historical and contemporary evidence for arachnoid villi, perineural routes, and dural lymphatics. Cell. Mol. Life Sci. 78, 2429–2457 (2021).

55. Lee, H. et al. Quantitative gd-dota uptake from cerebrospinal fluid into rat brain using 3d vfa-spgr at 9.4 t. Magn. resonance medicine 79, 1568–1578 (2018).

56. Pardridge, W. M. Csf, blood-brain barrier, and brain drug delivery. Expert. opinion on drug delivery 13, 963–975 (2016).

57. Hornkjol, M. et al. Csf circulation and dispersion yield rapid clearance from intracranial compartments. bioRxiv (2022).

58. Chagnot, A., Barnes, S. R. & Montagne, A. Magnetic resonance imaging of blood–brain barrier permeability in dementia. Neuroscience 474, 14–29 (2021).

59. Cserr, H., Cooper, D., Suri, P. & Patlak, C. Efflux of radiolabeled polyethylene glycols and albumin from rat brain. Am. J. Physiol. Physiol. 240, F319–F328 (1981).

60. Bradbury, M., Cserr, H. & Westrop, R. Drainage of cerebral interstitial fluid into deep cervical lymph of the rabbit. Am. J. Physiol. Physiol. 240, F329–F336 (1981).

61. Szentistvanyi, I., Patlak, C. S., Ellis, R. A. & Cserr, H. F. Drainage of interstitial fluid from different regions of rat brain. Am. J. Physiol. Physiol. 246, F835–F844 (1984).

62. Rosenberg, G., Kyner, W. & Estrada, E. Bulk flow of brain interstitial fluid under normal and hyperosmolar conditions. Am. J. Physiol. Physiol. 238, F42–F49 (1980).

63. Ray, L., Iliff, J. J. & Heys, J. J. Analysis of convective and diffusive transport in the brain interstitium. Fluids Barriers CNS 16, 1–18 (2019).

64. Bulat, M. & Klarica, M. Recent insights into a new hydrodynamics of the cerebrospinal fluid. Brain research reviews 65, 99–112 (2011).

65. Myung, J. et al. The choroid plexus is an important circadian clock component. Nat. communications 9, 1–13 (2018).

66. Hablitz, L. M. et al. Circadian control of brain glymphatic and lymphatic fluid flow. Nat. communications 11, 1–11 (2020).

67. Bojarskaite, L. et al. Sleep cycle-dependent vascular dynamics enhance perivascular cerebrospinal fluid flow and solute transport. bioRxiv (2022).

68. Fultz, N. E. et al. Coupled electrophysiological, hemodynamic, and cerebrospinal fluid oscillations in human sleep. Science 366, 628–631 (2019).

69. Glowinski, R., Song, Y., Yuan, X. & Yue, H. Bilinear optimal control of an advection-reaction-diffusion system. SIAM Rev. 64, 392–421 (2022).

70. Dembitskaya, Y. et al. Shadow imaging for panoptical visualization of brain tissue in vivo. Res. Gate (2022).

71. Fischl, B. Freesurfer. Neuroimage 62, 774–781 (2012).

72. Mardal, K.-A., Rognes, M. E., Thompson, T. B. & Valnes, L. M. Mathematical modeling of the human brain: From magnetic resonance images to finite element simulation (2022).

73. Troyetsky, D. E., Tithof, J., Thomas, J. H. & Kelley, D. H. Dispersion as a waste clearance mechanism in pressure-driven flow through open penetrating perivascular spaces. 73rd Annu. Meet. APS Div. Fluid Dyn. (2020).

74. Alnæs, M. et al. The fenics project version 1.5. Arch. Numer. Softw. 3 (2015).

75. Evans, L. C. Partial differential equations, vol. 19 (American Mathematical Soc., 2010).

76. Liu, D. C. & Nocedal, J. On the limited memory bfgs method for large scale optimization. Math. programming 45, 503–528 (1989).

77. Farrell, P. E., Ham, D. A., Funke, S. W. & Rognes, M. E. Automated derivation of the adjoint of high-level transient finite element programs. SIAM J. on Sci. Comput. 35, C369–C393 (2013).

78. Virtanen, P. et al. Scipy 1.0: fundamental algorithms for scientific computing in python. Nat. methods 17, 261–272 (2020).

