## Supplementary material for "Human brain solute transport quantified by glymphatic MRI-informed biophysics during sleep and sleep deprivation"

DRAFT

### S1. Evaluation of multi-modal data integration choices

The original clinical study (1), which data the present study rely on, involved 24 subjects. All subjects had T1-weighted MR images taken at multiple time points after intrathecal administration. 15 out of the 24 subjects had $T_1$ maps and 18 subjects had associated diffusion tensor images (DTI) taken before injection of tracers. Since not all data were available for all subjects, we first investigate whether raw, filtered or averaged $T_1$ maps and DTI data would yield quantitatively different conclusions in Section S1.1. Second, we assess the effect of using subject-specific DTI versus averaged diffusion coefficients on forward simulations, in Section S1.2. Finally, transport parameters that were obtained from PDE-constrained optimization are assessed with both subject-specific and study-averaged $T_1$ and DTI to confirm that parameters do not differ substantially between methods (Tables S6 and S7).

**S1.1. Mapping signal intensities to concentrations .** The $T_1$ maps enable conversion of contrast-enhanced MR signal intensities $S$ (arbitrary unit) to tracer concentrations $c$ (mmol/L). Specifically, for each subject, time of scan $t > 0$, and voxel $a_i$, we stipulate that:

$$c_{\mathrm{mri}}(a_i, t) = \frac{1}{r_1}\left(\frac{1}{T_1(a_i, t)} - \frac{1}{T_1^0(a_i)}\right), \quad \text{[S1]}$$

where $T_1^0$ is the longitudinal relaxation time prior to tracer injection as given by the $T_1$ maps, $r_1$ is a constant depending on the contrast agent, and $T_1(a_i, t)$ are the longitudinal relaxation times after tracer injection (2). In agreement with (1), we normalize the contrast-enhanced signal intensities and compute

$$T_1(a_i, t) = f^{-1}\left(\frac{S(a_i, t)}{S_0(a_i)} f(T_1^0)\right), \quad \text{[S2]}$$

where $S_0$ is the pre-contrast $T_1$ weighted MRI signal and $f$ is a monotone, nonlinear function specific to the MRI sequence used (3). Parameters for the $T_1$ map acquisition using the MOLLI5(3)3 (4) sequence, and hence the function $f$, are as previously established (3).

For 15 out of the 24 subjects in the present study, $T_1$ map images are available. The $T_1$ values are $900\,\mathrm{ms} \lesssim T_1 \lesssim 1400\,\mathrm{ms}$ in the parenchyma. Since the $T_1$ maps have lower resolution ($176 \times 176 \times 71$ voxels compared to $256 \times 256 \times 256$ in $T_1$ weighted images), significantly differing voxel value range, and are not available for all the participants, we investigate the importance of subject-specific $T_1$ map images for accurate quantification of tracer concentration in the brain. We consider three choices of $T_1$ maps for calculating the subject-specific concentrations, 1) using raw $T_1$ map, 2) using filtered $T_1$ map, and 3) using a group averaged $T_1$ map image. In detail, for the second method, for every of these 15 subjects, we create median-filtered $T_1$ maps by assigning the median $T_1$ map value of every subregion $\Omega_i$ (around 180 regions) as defined in the FreeSurfer parcellation file (wmparc.mgz) (5) to that region,

$$T_1(a_i) = \mathrm{median}_{y \in \Omega_i} T_1(y) \tag{S3}$$

where $\Omega_i$ indicates the FreeSurfer subregion that includes the voxel $a_i$. Finally, for the third method, we compute the average of the 15 available subjects as

$$\overline{T}_{1,k} = \frac{1}{15} \sum_{j=1}^{15} \frac{1}{\Omega_k^j} \sum_{a_i \in \Omega_k^j} T_1(a_i) \tag{S4}$$

where the index $k \in \{g, w, b\}$ indicates three brain subregions $\Omega_k$ cerebral cortex, subcortical subcortical white matter and the brain stem. This yields

$$\overline{T}_1(a_i) = \begin{cases} 1002\,\mathrm{ms} & \text{for } a_i \in \text{ subcortical white matter} \\ 1274\,\mathrm{ms} & \text{for } a_i \in \text{ cerebral cortex} \\ 1127\,\mathrm{ms} & \text{for } a_i \in \text{ brain stem} \end{cases} \tag{S5}$$

Using these mean values, as well as the filtered values Eq. (S3) and raw $T_1$ maps, we then estimate the concentration in every voxel $a_i$ using Eq. (S1) and Eq. (S2) with $\overline{T}_1$. The voxel-based concentration values $c_{\mathrm{mri}}(a_i, t)$ are then used to represent the brain-wide concentration field $c(x, t)$ on the computational mesh using continuous linear Lagrange elements. The total amount of tracer imaged in the brain at time $t$ is computed as

$$\int_\Omega c(x, t)\,\mathrm{d}x \tag{S6}$$

and the total amount of tracer imaged on the brain surface at time $t$, is computed as

$$\int_{\partial\Omega} c(x, t)\,\mathrm{d}S. \tag{S7}$$

DRAFT

The resulting concentration values in the brain and on the surface are displayed in Figs. S1A-B. The three $T_1$ maps yield average brain concentration around 0.1 mmol/L after 6 and 24 hours, and there are no significant differences between the choices neither in concentration in the brain nor at the surface (at all time points). The estimates differ the most at 24 h; in the brain the concentration differs by $7 \pm 2\,\%$ between method 1 and 2, and $11 \pm 4\,\%$ by 2 and 3. At the pial surface, the values differ by $13 \pm 3\,\%$ between method 1 and 2, and $24 \pm 7\,\%$ by 2 and 3. Fig. S1C compares simulation and data for the three choices of $T_1$ map. The conclusion drawn from either choice is the same; simulations underestimate the influx at $\sim 6\,\mathrm{h}$ by $59, 61, 58\,\%$ and overestimate what remains in the brain after 48 h by $81, 93, 85\,\%$ for the $T_1$ maps 1-3, respectively.

Across the subjects, the resulting concentrations correlate strongly with the MRI signal increase inside the brain (0.97-0.99) and at the pial surface (0.89-0.99), cf. Fig. S1D-K. Further, the concentrations obtained from averaged $T_1$ maps increase correlate strongly with those obtained by raw $T_1$ map, (0.98-1.00) for brain-concentrations and (0.97-0.99) at the pial surface, cf. Fig. S1L-S.

These observations justify the use of synthetic $T_1$ maps $\overline{T}_1$ to estimate the concentration in the 9 / 24 subjects where no $T_1$ map is available. We hence use method 3 (group averaged $T_1$ map image) to estimate the concentration in all 24 subjects.

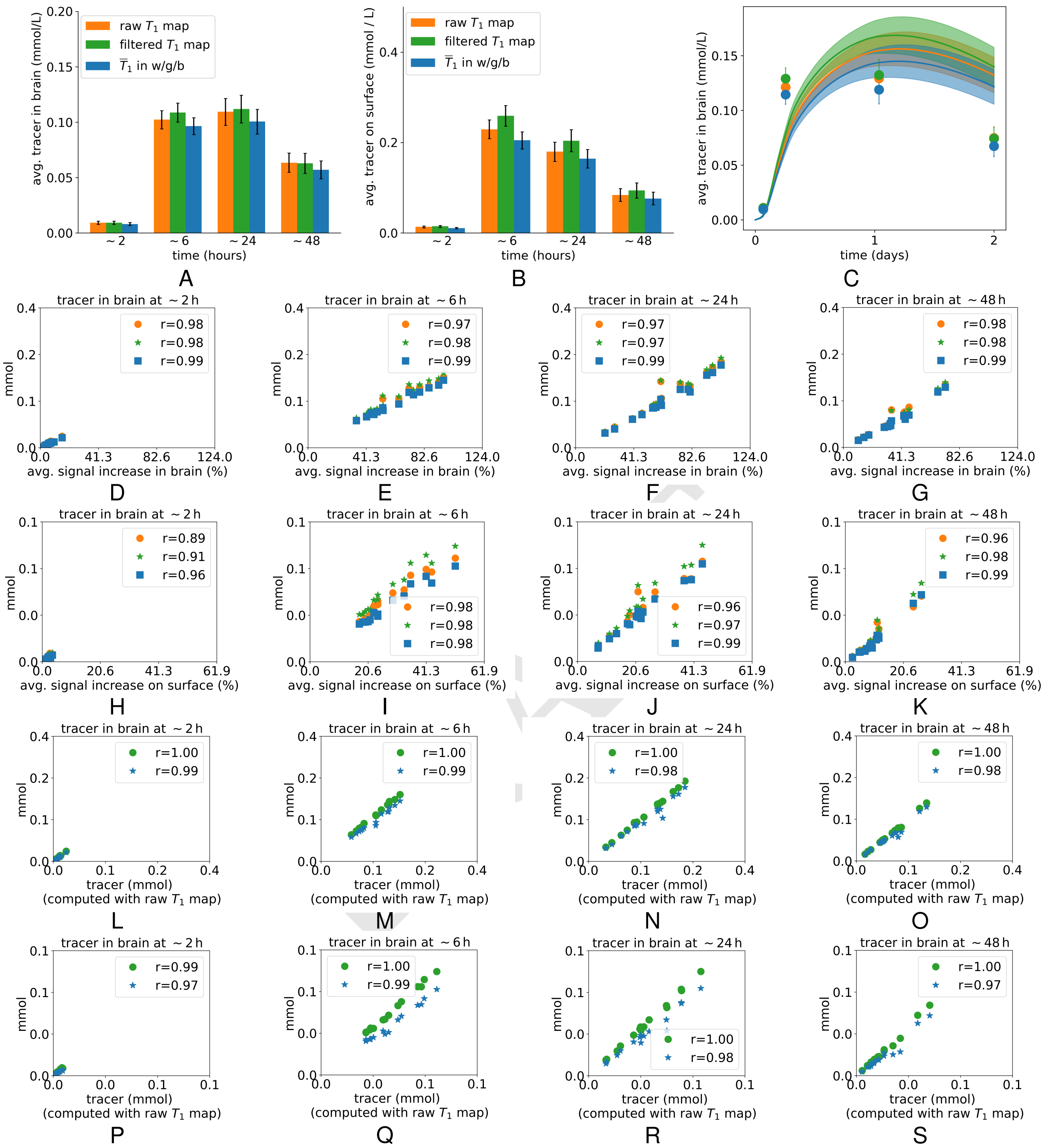

**Fig. S1.** Tracer in brain (A) and at the pial surface (B) as computed with the three different choices for $T_1$ maps. (C) Measured (dots) and simulated (continuous lines) tracer in the brain, averaged over the $n = 15$ subjects where good $T_1$ maps are available. Error bars and shaded regions indicate standard error of the mean. (D-G) correlation between signal increase and tracer in the brain for different time points. (H-K) correlation between signal increase and tracer at the pial surface for different time points. (L-O) correlation between tracer as computed with raw $T_1$ map and computed with the other two methods, in the brain, for different time points. (P-S) correlation between tracer as computed with raw $T_1$ map and computed with the other two methods, at the pial surface, for different time points.

**S1.2. Influence of using diffusion tensor images on simulation results .** From the DTI scans, we directly obtain subject-specific voxel-wise apparent diffusion tensors for water $\mathbf{D_0^*}$. The apparent diffusion coefficient for water $D_0^*$ is computed as the mean diffusivity:

$$D_0^* = \frac{1}{3}\mathrm{tr}(\mathbf{D_0^*}).$$

Given also the free diffusion coefficient of water $D_0$ we estimate the tortuosity of the brain as $\lambda^2 = D_0/D_0^*$, and (assuming the tortuosity for water and CSF tracer is the same) the apparent diffusion coefficient of the CSF tracer is thus given by:

$$D^* = \frac{D}{\lambda^2} = \frac{D}{D_0}D_0^*,$$

where $D = 3.8 \times 10^{-4}\,\mathrm{mm^2 s^{-1}}$ is the free diffusion coefficient of the CSF tracer gadobutrol (3) used in this study. To convert diffusion tensors from water to CSF tracer diffusivity, we correspondingly scale the DTI values by the ratio $D/D_0$ to obtain the diffusion tensor in each voxel $a_i$:

$$\mathbf{D}^*(a_i) = \frac{D}{D_0}\mathbf{D_0^*}(a_i). \qquad \text{[S8]}$$

For 18 out of 24 subjects, DTI taken at baseline (before tracer injection) were available. We first checked if the mean diffusivities estimated from these images differs between the sleep and sleep deprivation group. From the diffusion tensor $\mathbf{D}_0^*$ for water we computed the averaged mean diffusivity $\int \frac{1}{3}\mathrm{tr}(\mathbf{D}_0^*)\,\mathrm{d}x$ in subcortical white matter and the cerebral cortex for every subject. The values are illustrated in Fig. S2. No significant differences between groups were found in either subcortical white matter or the cerebral cortex.

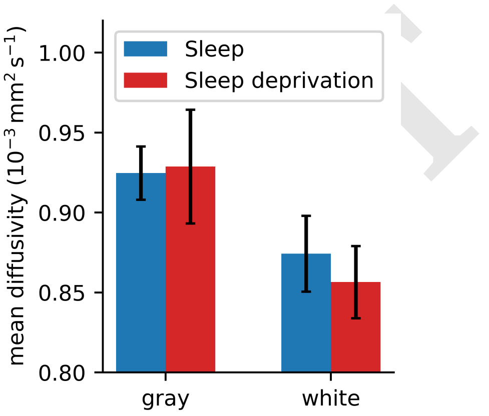

**Fig. S2.** The average water diffusivity in subcortical white matter and the cerebral cortex does not vary significantly between the two subject groups. Error bars represent standard error of the group mean.

For 6 out of the 24 subjects, no DTI was available. In order to incorporate these subjects into the analysis, we investigated the feasibility of using mean scalar diffusion coefficients instead of subject-specific DTI for the diffusivity $D^*$ in Eq. (1). From the $j = 1, \ldots, 18$ subjects where DTI was available, we computed study average mean scalar diffusion coefficients as

$$\overline{D}_i = \frac{1}{18}\sum_{j=1}^{18}\frac{1}{\Omega_i^j}\int_{\Omega_i^j}\frac{1}{3}\mathrm{tr}(\mathbf{D}_0^*)\,\mathrm{d}x \qquad \text{[S9]}$$

where the index $i \in \{g, w, b\}$ indicates three brain subregions $\Omega_i^j$ cerebral cortex, subcortical white matter matter and the brain stem of subject $j$. Using these values, we then solve Eq. (1) with $D$ as defined in Eq. (S9),

$$D(x) = \begin{cases} \overline{D}_g = 1.17 \times 10^{-4}\,\mathrm{mm^2 s^{-1}} & \text{for } x \in \text{ cerebral cortex} \\ \overline{D}_w = 1.10 \times 10^{-4}\,\mathrm{mm^2 s^{-1}} & \text{for } x \in \text{ subcortical white matter matter} \\ \overline{D}_b = 1.34 \times 10^{-4}\,\mathrm{mm^2 s^{-1}} & \text{for } x \in \text{ brain stem} \end{cases}$$

as well as the spatially varying diffusion tensor Eq. (S8). The resulting simulated concentration distributions after 48 h are shown for an example subject in Fig. S3. Both choices lead to visually identical predictions, and in the shown example, the relative $l^1$-difference is 7 %.

Fig. S4A illustrates the tracer concentration over time as simulated with both choices averaged over the 18 subjects with DTI. The relative difference between the curves is about 1 %. In both cases, the amount of simulated tracer after 5-7 h is about 60 % too small as compared to data. After two days it is 200 % too large in both choices.

Figs. S4B-D further show the correlation between tracer in the brain as measured and simulated using DTI/simulated with averaged $D$ at 6, 24, 48 hours. Overall, the correlation is in the range (0.91-0.95). At every time point, the correlation measures for both methods are nearly identical.

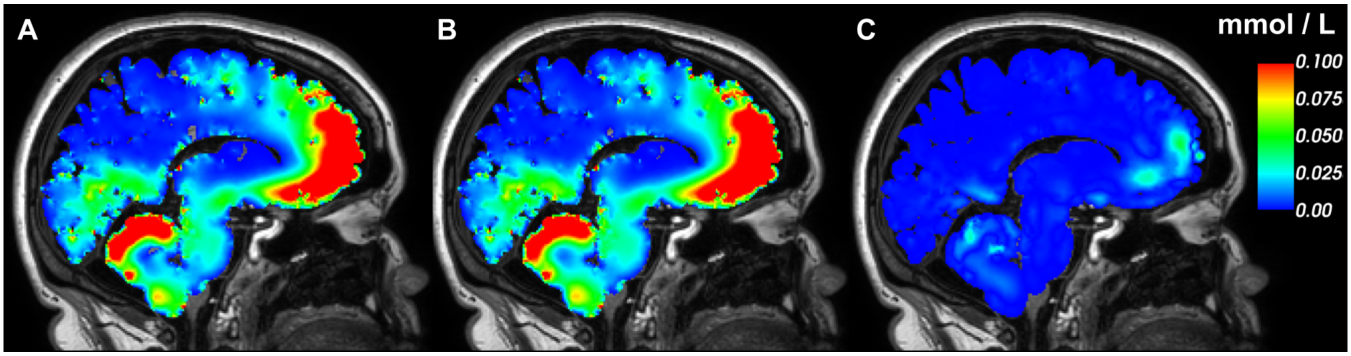

**Fig. S3.** Simulated tracer distribution after 48 hours in example subject 14, using DTI (A) and averaged diffusion coefficients (B). The absolute error between A-B is shown in (C) in the same color scale.

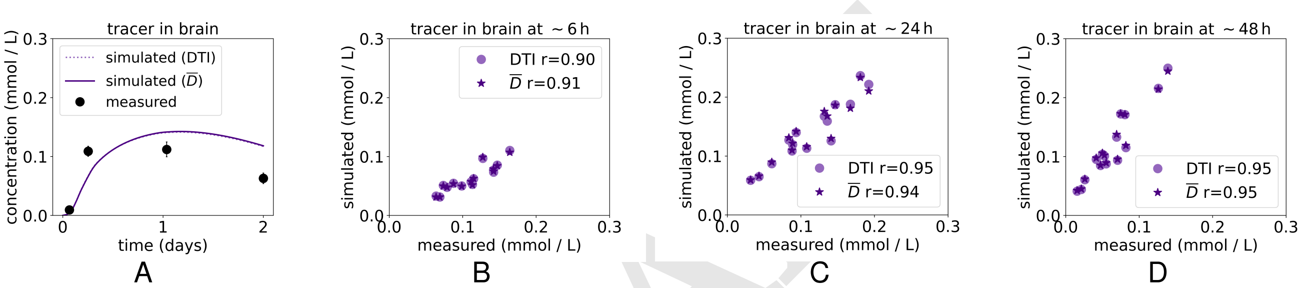

**Fig. S4.** A. Tracer in brain as measured (dots) and simulated with DTI and avg. $\overline{D}$ (continuous lines) over two days. B-D. Correlation between tracer in the brain as measured (horizontal axis) and simulated using DTI / simulated with averaged $D$ (vertical axis) at 6, 24, 48 hours for the subjects where DTI is available.

In summary, the difference in simulated averaged concentrations between choosing an average diffusion constant versus using the diffusion tensor from DTI differed by less than 1 % (Fig. S4a). Furthermore, the correlation coefficients between the two methods were all $r = 1.00$ at 6, 24 and 48 hours. Based on these results, in each of the 24 subject we choose to assign the mean diffusion coefficients given by Eq. (S10) to a given region (cerebral cortex, subcortical white matter matter and brain stem).

### S2. Numerics and computation

**S2.1. Computational mesh characteristics.** For each subject, left and right hemisphere pial surfaces, left and right cerebral cortex/subcortical white matter interface surfaces as well as ventricular surfaces were extracted from the T1-weighted MR images prior to tracer injection using FreeSurfer (6). The surfaces were refined and smoothened using computational geometry techniques prior to construction of volumetric meshes of the brain parenchyma at low-, standard- and high resolution via SVMTK (7). Table S1 gives an overview of the complexity and approximability of the meshes at different resolution levels.

| subject ID | Low-res | | | | Standard | | | | High-res | | | |
|---|---|---|---|---|---|---|---|---|---|---|---|---|
| | $n_v$ | $n_c$ | $h_{\min}$ | $h_{\max}$ | $n_v$ | $n_c$ | $h_{\min}$ | $h_{\max}$ | $n_v$ | $n_c$ | $h_{\min}$ | $h_{\max}$ |
| 1 | 56663 | 268451 | 0.21 | 10.42 | 205649 | 917870 | 0.10 | 5.23 | | | | |
| 2 | 75080 | 374297 | 0.40 | 10.70 | 213735 | 1012961 | 0.31 | 5.41 | | | | |
| 3 | 73549 | 385438 | 1.05 | 10.88 | 247863 | 1167963 | 0.52 | 5.47 | 9456 | 49843 | 2.42 | 23.81 |
| 4 | 76920 | 392357 | 1.01 | 10.99 | 229201 | 1085103 | 0.52 | 5.51 | 864112 | 4044782 | 0.25 | 2.77 |
| 5 | 78000 | 394201 | 1.10 | 11.02 | 229561 | 1085115 | 0.43 | 5.55 | | | | |
| 6 | 74747 | 404183 | 1.06 | 10.66 | 250598 | 1205522 | 0.45 | 5.37 | | | | |
| 7 | 80552 | 421045 | 0.94 | 11.01 | 250010 | 1192453 | 0.09 | 5.43 | 971769 | 4545533 | 0.13 | 2.73 |
| 8 | 73893 | 376981 | 0.96 | 10.30 | 236044 | 1103313 | 0.45 | 5.18 | 925360 | 4327308 | 0.24 | 2.60 |
| 9 | 74305 | 380830 | 0.89 | 10.26 | 230342 | 1086651 | 0.50 | 5.14 | 899409 | 4205827 | 0.15 | 2.58 |
| 10 | 79983 | 403220 | 0.98 | 10.59 | 228810 | 1080081 | 0.43 | 5.38 | 855631 | 4023766 | 0.23 | 2.69 |
| 11 | 80824 | 404862 | 0.60 | 10.81 | 228540 | 1075838 | 0.52 | 5.42 | 869291 | 4044744 | 0.10 | 2.73 |
| 12 | 86192 | 448242 | 0.87 | 10.95 | 256069 | 1225482 | 0.48 | 5.49 | 973581 | 4554988 | 0.24 | 2.75 |
| 13 | 81574 | 424664 | 1.00 | 10.83 | 253784 | 1199128 | 0.51 | 5.43 | 955595 | 4428214 | 0.04 | 2.72 |
| 14 | 81143 | 423197 | 1.02 | 10.91 | 240837 | 1167721 | 0.50 | 5.47 | 950203 | 4453739 | 0.24 | 2.75 |
| 15 | 79428 | 406316 | 1.01 | 10.90 | 234481 | 1117785 | 0.54 | 5.58 | 903415 | 4243049 | 0.23 | 2.77 |
| 16 | 79059 | 405030 | 0.94 | 10.69 | 239084 | 1136336 | 0.51 | 5.40 | 915324 | 4314036 | 0.02 | 2.70 |
| 17 | 80163 | 415968 | 1.04 | 11.06 | 239221 | 1141950 | 0.55 | 5.60 | 925175 | 4293556 | 0.03 | 2.81 |
| 18 | 78527 | 391206 | 0.95 | 11.37 | 211243 | 1006364 | 0.47 | 5.73 | 766404 | 3613485 | 0.24 | 2.87 |
| 19 | 78125 | 400132 | 1.04 | 10.67 | 238326 | 1129354 | 0.47 | 5.35 | 926311 | 4339819 | 0.16 | 2.68 |
| 20 | 76058 | 379266 | 1.10 | 10.46 | 220072 | 1038529 | 0.50 | 5.31 | 829541 | 3944186 | 0.23 | 2.66 |
| 21 | 82373 | 420395 | 1.00 | 10.40 | 251776 | 1188577 | 0.47 | 5.22 | 954926 | 4500535 | 0.22 | 2.61 |
| 22 | 82737 | 425841 | 0.96 | 10.79 | 250220 | 1184843 | 0.48 | 5.43 | 1039209 | 4761980 | 0.23 | 2.72 |
| 23 | 81636 | 404870 | 0.97 | 10.62 | 234802 | 1104395 | 0.47 | 5.35 | 882364 | 4163055 | 0.22 | 2.70 |
| 24 | 75769 | 390828 | 0.95 | 10.37 | 240209 | 1140383 | 0.46 | 5.19 | | | | |

**Supplementary Table S1. Overview of computational mesh characteristics for each subject for low- standard and high-resolution meshes. Here, $n_v$ denotes the number of mesh vertices, $n_c$ the number of mesh cells, and the minimal and maximal mesh cell sizes $h_{\min}$ and $h_{\max}$ (in mm) are computed as the minimal and maximal mesh cell circumradius $\times 2$, respectively.**

DRAFT

**S2.2. Numerical verification and convergence of the forward problem .** In order to access the numerical convergence of the forward diffusion simulations, we performed simulations of Eq. (1) using different mesh resolutions and time step sizes. In detail, for two example subjects, we solve Eq. (1) over a time span of two days with $\alpha = 1, r = 0, \phi = 0$ with time steps of 30, 20, 10 min and meshes of resolution parameter 64, 32, 16 (corresponding to maximal mesh cell sizes of $\sim 3, 5, 11$ mm, respectively).

Fig. S5 shows the simulated brain-wide CSF tracer concentration over a time span of two days for the two example subjects. The low resolution meshes yield visibly lower concentrations compared to standard and high resolution meshes; integrated concentrations differ to a relative error of 14 % and 20 % between low and high resolution meshes for example subject 4 and 14, respectively. However, standard and high resolution meshes yield comparable results, the simulated concentrations differ to a relative error of 1 % and 4 % for example subject 4 and 14, respectively.

The errors between simulated tracer with low and high resolution meshes and time step of 20 min are 15 % after 48 hours for subject 4, and 19 % after 48 hours for subject 14. For standard and high resolution meshes, the errors between simulated tracer after 48 hours are 1 % after 48 hours for subject 4, and 3 % for subject 14, indicating that the conclusion drawn from either standard or high resolution mesh is the same.

The data shown in Table S2 show that the simulated concentration after 48 h agrees up to the third digit for standard and high resolution meshes and all time step sizes. We hence conclude that standard resolution meshes yield accurate results and use meshes of this resolution to produce the results reported in the main manuscript. For the inverse velocity identification we perform simulations with low resolution meshes in addition to investigate the convergence of the method, cf. Section S3.1. For the forward diffusion simulations with $\alpha \in \{1, 2, 3, 4, 5\}$ and $r = \phi = 0$, we use a time step size of 20 min to limit the computational cost.For the inverse identification of parameters in the enhanced diffusion–local clearance model, we assess the influence of time step size on the results separately in Section S4.3.

**Supplementary Table S2. Brain-wide CSF tracer concentration (mmol / L) at 48 hours, data and diffusion simulation $(\alpha = 1, r = 0, \phi = 0)$, for different time/mesh resolutions for two example subjects.**

| | $dt$ | $n = 16$ | $n = 32$ | $n = 64$ |
|---|---|---|---|---|
| data | - | 0.046 | 0.046 | 0.046 |
| simulation | 30 | 0.101 | 0.118 | 0.119 |
| | 20 | 0.101 | 0.118 | 0.119 |
| | 10 | 0.100 | 0.117 | 0.118 |

**(a) Example subject 4**

| | $dt$ | $n = 16$ | $n = 32$ | $n = 64$ |
|---|---|---|---|---|
| data | - | 0.014 | 0.014 | 0.013 |
| simulation | 30 | 0.037 | 0.045 | 0.046 |
| | 20 | 0.037 | 0.045 | 0.046 |
| | 10 | 0.037 | 0.045 | 0.046 |

**(b) Example subject 14**

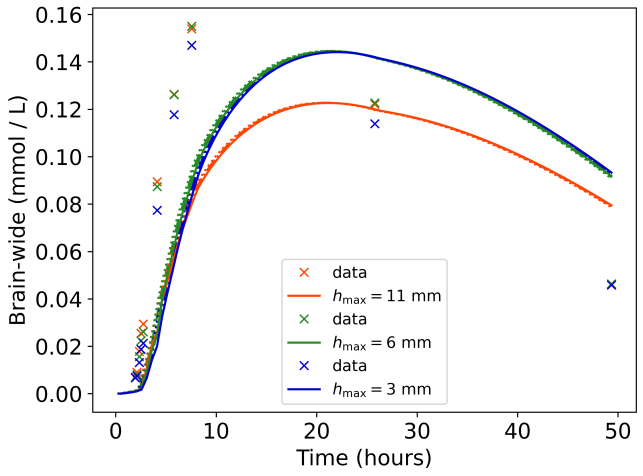

**(a)** Example subject 4

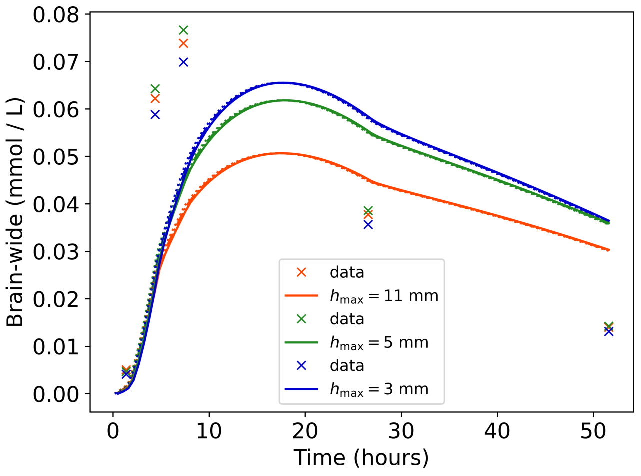

**(b)** Example subject 14

**Fig. S5.** Dependence of the tracer concentration obtained by solving Eq. (1) with $\alpha = 1, r = 0, \phi = 0$ using meshes of different resolution. The different time step sizes of 30, 20 and 10 min yield similar results and are encoded by line styles "-", ".-" and "..", respectively.

### S3. Additional information for the inverse velocity field identification

This section gives additional methodological details and results for the inverse estimation of the velocity field $\phi$ in the following special case of Eq. (1) (with $\alpha = 1, r = 0$):

$$\frac{\partial c}{\partial t} - \nabla \cdot D^* \nabla c + \nabla \cdot (c\phi) = 0 \quad \text{in } \Omega, \quad t > t_0. \tag{S10}$$

**S3.1. Numerical verification of the inverse velocity identification.** In order to evaluate the numerical accuracy of the computationally estimated velocity fields, we considered a series of experiments for different computational mesh resolutions (low and standard resolutions) and different regularization parameters ($\beta \in \{10^{-3}, 10^{-4}, 10^{-5}\}$) for each subject and for each of the time intervals 0–6h, 6–24h and 24–48h. For each of these simulations, we first evaluated the robustness and effectiveness of the optimization algorithm by comparing the value of the initial functional $J_0$ and final (optimized) functional $J_N$, the maximal component of the optimized velocity field $\max \phi$ and its relative differential from its penultimate value (see Table S3 for the time interval 24–48h). Two subjects (3, 23) were missing contrast MR scans in the ∼6h time interval, while three subjects (1, 2, 5) where missing scans in the 24–48h interval. These subjects were therefore excluded from further analysis in the relevant time interval.

For the ∼0–6h intervals, the objective functionals were substantially reduced for all remaining subjects and numerical parameters ($n$, $\beta$), typically by two–three orders of magnitude (by a factor ranging from 6 to 614), and the relative differential in the maximal velocity component was less than 1.84% for all cases. Similarly, for the ∼6–24h intervals, all optimizations were considered satisfactory with objective functional reductions of $26 - 4765\times$ and a maximal relative differential of 1.06% (data not shown). For the ∼24-48h intervals, the optimization algorithm did not converge satisfactorily for subject IDs 6, 12 and 14 for any $\beta$, n (little to no reduction in the objective functionals). For these artefactual cases, the maximal velocity component estimates also varied substantially with $\beta$. We therefore excluded the numerical results also from these subjects for the interval 24–48h from the subsequent analyses. For the remaining subjects, the maximal relative differential in the maximal velocity component was 2.14% and the relative reduction in the objective functional substantial (8–2098×).

We next proceeded to evaluate the numerical accuracy of the optimal velocity estimates, in particular, we are interested in any dependence of the numerical results on the primary numerical parameters such as the mesh resolution $n$ and choice of regularization parameter $\beta$. For each numerical simulation (having passed the optimization algorithm evaluation described above), we computed the average velocity magnitude $\bar{\phi}$ for each velocity field $\phi = \phi(x) = (\phi_1(x), \phi_2(x), \phi_3(x))$ as

$$\bar{\phi} = |\Omega|^{-1} \int_\Omega |\phi| \,\mathrm{d}x, \quad |\Omega| = \int_\Omega 1 \,\mathrm{d}x, \quad |\phi|^2 = \phi_1^2 + \phi_2^2 + \phi_3^2,$$

as well as the $L^2$-norm of the associated estimated concentration field $c = c(x, t_1)$ solving Eq. (1) on the relevant time interval $(t_0, t_1)$, the $L^2$-norm of the corresponding target concentration $c_1$, and the relative difference between these values $\Delta t$. Based on the following analysis of these values, we use $n = 32$ and $\beta = 10^{-4}$ for the numerical simulations reported in the main text.

For the first time interval, ∼0–6h (data not shown), we observe that the relative difference in concentration norms is small for all experiments (typically less than 1%, with a maximal relative difference when $\beta = 10^{-4}$ of 3.84%). Moreover, we observe that the estimated average velocity shows only small variations (if any) with respect to the regularization parameter $\beta$ especially for the standard resolution meshes $n = 32$. However, we do observe some variations with respect to the mesh size $n$, indicating

DRAFT

| ID, t0 | $n$ | $J_0$ | $J_N$ | $\max \phi_N$ ($\Delta\phi$%) | $J_0$ | $J_N$ | $\max \phi_N$ ($\Delta\phi$ %) | $J_0$ | $J_N$ | $\max \phi_N$ ($\Delta\phi$ %) |
|---|---|---|---|---|---|---|---|---|---|---|
| 3, 24h | 16 | 1482.9 | 146.52 | 3.27 (0.44) | 1482.9 | 65.39 | 3.79 (0.18) | 1482.9 | 39.86 | 4.28 (0.10) |
| | 32 | 1450.4 | 143.04 | 2.24 (0.53) | 1450.4 | 73.71 | 6.17 (0.03) | 1450.4 | 54.00 | 8.05 (0.40) |
| 4, 24h | 16 | 5050.8 | 106.96 | 2.17 (1.53) | 5050.8 | 26.67 | 3.77 (0.20) | 5050.8 | 10.77 | 4.46 (0.15) |
| | 32 | 4805.5 | 99.86 | 2.71 (2.14) | 4805.5 | 29.48 | 5.99 (0.13) | 4805.5 | 12.13 | 6.10 (0.03) |
| 6, 24h | 16 | 5972.8 | 5889.23 | 4.47 (0.01) | 5972.8 | 5463.50 | 20.12 (0.00) | 5972.8 | 4271.71 | 68.19 (0.00) |
| | 32 | 5854.9 | 5744.12 | 6.19 (0.01) | 5854.9 | 5226.90 | 22.82 (0.02) | 5854.9 | 3863.81 | 77.73 (0.00) |
| 7, 24h | 16 | 1140.8 | 74.63 | 1.68 (0.42) | 1140.8 | 27.08 | 2.32 (0.09) | 1140.8 | 13.56 | 2.59 (0.32) |
| | 32 | 1156.6 | 81.85 | 3.62 (0.87) | 1156.6 | 37.48 | 7.11 (0.18) | 1156.6 | 33.49 | 8.01 (0.35) |
| 8, 24h | 16 | 12433.2 | 321.34 | 4.97 (0.20) | 12433.2 | 213.57 | 6.77 (0.16) | 12433.2 | 173.93 | 5.71 (0.44) |
| | 32 | 11553.1 | 313.16 | 4.67 (0.45) | 11553.1 | 218.26 | 4.88 (0.00) | 11553.1 | 224.74 | 4.98 (0.13) |
| 9, 24h | 16 | 1168.5 | 22.79 | 1.00 (0.40) | 1168.5 | 3.82 | 1.33 (0.03) | 1168.5 | 0.83 | 1.40 (0.05) |
| | 32 | 1057.9 | 23.84 | 1.94 (0.44) | 1057.9 | 5.69 | 2.53 (0.11) | 1057.9 | 3.79 | 2.59 (0.10) |
| 10, 24h | 16 | 7603.0 | 128.51 | 3.73 (0.57) | 7603.0 | 61.45 | 4.60 (0.13) | 7603.0 | 41.08 | 4.52 (0.41) |
| | 32 | 7091.7 | 120.73 | 1.91 (0.02) | 7091.7 | 65.64 | 2.32 (0.02) | 7091.7 | 63.68 | 2.78 (0.41) |
| 11, 24h | 16 | 3754.2 | 92.43 | 2.12 (0.64) | 3754.2 | 39.56 | 2.79 (1.08) | 3754.2 | 33.95 | 2.79 (0.10) |
| | 32 | 3489.3 | 217.79 | 1.97 (0.72) | 3489.3 | 74.39 | 2.48 (0.19) | 3489.3 | 61.79 | 2.40 (0.21) |
| 12, 24h | 16 | 1862.2 | 1462.40 | 6.59 (0.02) | 1862.2 | 942.03 | 19.23 (0.00) | 1862.2 | 428.08 | 39.68 (0.11) |
| | 32 | 1770.7 | 1342.24 | 6.45 (0.00) | 1770.7 | 830.80 | 18.42 (0.07) | 1770.7 | 354.04 | 52.17 (0.20) |
| 13, 24h | 16 | 2036.9 | 36.11 | 1.08 (0.40) | 2036.9 | 11.49 | 1.31 (0.07) | 2036.9 | 8.32 | 1.42 (0.74) |
| | 32 | 1826.9 | 37.23 | 1.06 (0.02) | 1826.9 | 17.77 | 1.26 (0.18) | 1826.9 | 15.75 | 1.26 (0.27) |
| 14, 24h | 16 | 1064.1 | 631.28 | 5.82 (0.00) | 1064.1 | 337.30 | 14.60 (0.01) | 1064.1 | 125.84 | 23.48 (0.10) |
| | 32 | 1185.6 | 634.80 | 6.72 (0.01) | 1185.6 | 310.04 | 15.28 (0.03) | 1185.6 | 107.10 | 27.69 (0.49) |
| 15, 24h | 16 | 432.1 | 14.77 | 0.94 (0.84) | 432.1 | 5.30 | 1.56 (0.14) | 432.1 | 3.05 | 1.88 (0.30) |
| | 32 | 421.3 | 15.41 | 1.30 (0.11) | 421.3 | 6.78 | 1.51 (0.42) | 421.3 | 5.49 | 1.61 (0.46) |
| 16, 24h | 16 | 6243.6 | 223.14 | 3.77 (0.20) | 6243.6 | 160.44 | 4.53 (0.02) | 6243.6 | 131.29 | 4.56 (0.11) |
| | 32 | 6037.8 | 276.89 | 3.06 (0.02) | 6037.8 | 253.44 | 3.39 (0.37) | 6037.8 | 209.95 | 3.34 (0.10) |
| 17, 24h | 16 | 4513.1 | 48.04 | 1.13 (0.34) | 4513.1 | 12.15 | 1.37 (0.73) | 4513.1 | 7.01 | 1.42 (0.90) |
| | 32 | 3857.9 | 51.19 | 1.75 (0.18) | 3857.9 | 26.80 | 1.87 (0.32) | 3857.9 | 21.06 | 1.92 (0.17) |
| 18, 24h | 16 | 4136.5 | 252.19 | 4.67 (0.05) | 4136.5 | 51.14 | 10.35 (0.42) | 4136.5 | 9.77 | 12.90 (0.43) |
| | 32 | 4203.6 | 244.07 | 6.57 (0.70) | 4203.6 | 60.18 | 6.22 (0.12) | 4203.6 | 16.18 | 7.25 (0.36) |
| 19, 24h | 16 | 2051.3 | 25.86 | 0.99 (0.16) | 2051.3 | 7.11 | 1.08 (0.19) | 2051.3 | 4.51 | 1.09 (0.18) |
| | 32 | 1872.7 | 25.74 | 1.48 (0.26) | 1872.7 | 10.74 | 1.68 (0.07) | 1872.7 | 10.48 | 1.73 (0.00) |
| 20, 24h | 16 | 3222.5 | 39.09 | 1.25 (0.15) | 3222.5 | 8.31 | 2.05 (0.55) | 3222.5 | 2.86 | 2.73 (0.13) |
| | 32 | 3286.4 | 43.31 | 2.22 (1.59) | 3286.4 | 11.06 | 4.39 (0.01) | 3286.4 | 4.91 | 4.72 (0.07) |
| 21, 24h | 16 | 2526.6 | 32.42 | 1.30 (0.05) | 2526.6 | 9.98 | 1.41 (0.03) | 2526.6 | 8.24 | 1.41 (0.22) |
| | 32 | 2806.9 | 36.07 | 1.60 (0.07) | 2806.9 | 19.12 | 1.69 (0.07) | 2806.9 | 15.32 | 1.70 (0.13) |
| 22, 24h | 16 | 2260.7 | 50.58 | 1.94 (0.28) | 2260.7 | 17.68 | 2.93 (0.19) | 2260.7 | 12.66 | 3.23 (0.41) |
| | 32 | 2045.9 | 47.70 | 2.02 (0.59) | 2045.9 | 15.10 | 2.32 (0.70) | 2045.9 | 11.69 | 2.38 (0.82) |
| 23, 24h | 16 | 4397.1 | 18.09 | 1.25 (0.04) | 4397.1 | 3.89 | 1.24 (0.12) | 4397.1 | 2.10 | 1.24 (0.01) |
| | 32 | 4600.0 | 23.30 | 1.57 (0.89) | 4600.0 | 10.81 | 1.68 (0.53) | 4600.0 | 8.34 | 1.37 (0.24) |
| 24, 24h | 16 | 4742.9 | 29.20 | 1.60 (0.06) | 4742.9 | 7.58 | 1.72 (0.14) | 4742.9 | 4.90 | 1.78 (0.05) |
| | 32 | 4348.6 | 39.36 | 1.10 (0.37) | 4348.6 | 18.99 | 1.14 (0.37) | 4348.6 | 19.66 | 1.16 (0.23) |

**Supplementary Table S3. (24–48h) Key diagnostics for the optimization-based velocity estimation algorithm for low ($n = 16$) and standard ($n = 32$) resolution meshes, and three values of the regularization parameter $\beta \in \{10^{-3}, 10^{-4}, 10^{-5}\}$ (set of columns) for the time intervals 24–48h. $J_0$: value of the objective functional without convective velocity i.e. $\phi = 0$, $J_N$: final value of the objective functional after algorithmic completion, $\max \phi_N$ (mm/h): maximal component of the optimal velocity field, $\Delta\phi\% = |\max \phi_{N-1} - \max \phi_N| / \max \phi_N 100\%$: relative difference between maximal component of the penultimate and ultimate velocity optimization iterate (in percent); low relative difference indicates that the numerical algorithm has converged.**

DRAFT

that a comparison with a high resolution mesh could give further insights with regard to the numerical convergence of the method. For the time interval ∼6–24h (data not shown), we also observe low relative differences in concentration norms for all experiments: maximal relative difference when $\beta = 10^{-4}$ is 0.83%. The average velocity is highly robust with respect to $\beta$, and also its variation with respect to $n$ is relatively small.

For the time interval ∼24–48h (Table S4), we observe higher relative differences in concentration norms for $\beta = 10^{-3}$ (up to 10%), but the relative difference remains low for $\beta = 10^{-4}, 10^{-5}$. The maximal relative difference for $\beta = 10^{-4}$ is 1.63%. We observe some variation in the average velocity norm for $\beta = 10^{-3}$, while variation is less when comparing $\beta = 10^{-4}$ and $10^{-5}$. We observe some, but generally only small, variations between mesh sizes, and thus consider $\beta = 10^{-4}$ and $n = 32$ to yield more accurate numerical solutions.

**S3.2. Subject-level statistics of estimated convective flow field averages.** The subject-specific averaged velocity magnitudes, averaged brain-wide ($v$), over the cerebral cortex ($v_g$), over the subcortical white matter matter ($v_w$) and over the brain stem ($v_s$) are given in Table S5 for 6–24h (top) and 24–48h (bottom).

The average velocities in the intervals 6–24 h and 24–48h correlate only weakly with $r \approx 0.3$ in all regions of interest, cf. Fig. S6.

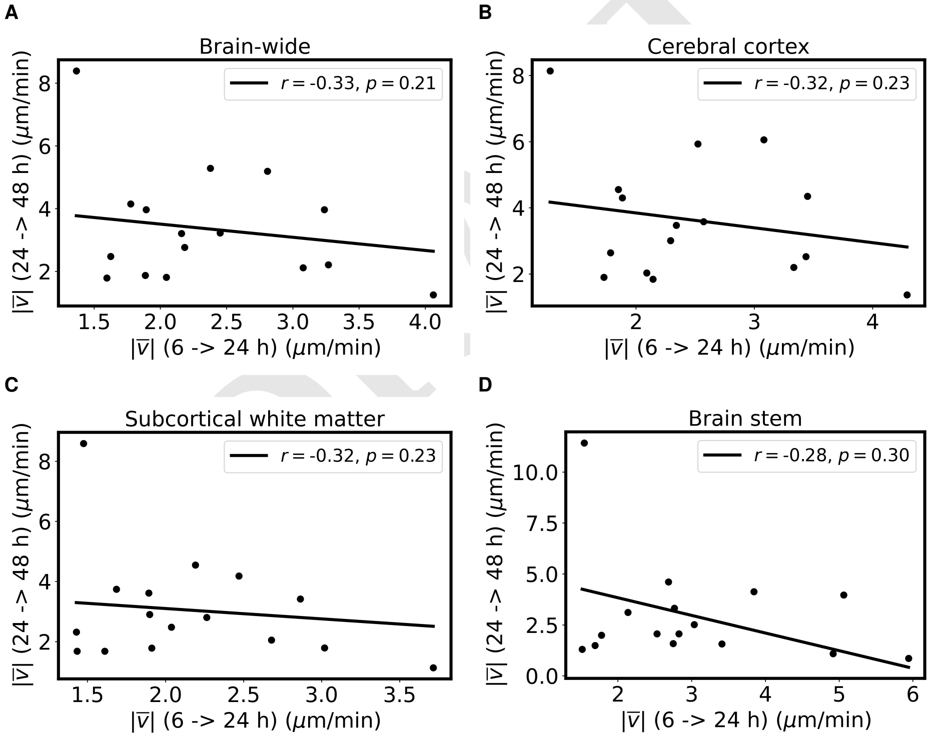

**Fig. S6.** Correlation between average velocities in 6-24 h interval to 24-48 h interval.

**S3.3. Influence of DTI and T1 map on identified velocity fields.** We further assessed how sensitive the identified velocity fields are with respect to data integration choices. Using (a) subject-patient specific diffusion tensors and $T_1$ maps and (b) study averaged constants for $D$ and $T_1$ for the 15 subjects where this data is available, we compare quantities of interest of the resulting velocity field obtained with regularization parameter $\beta = 10^{-4}$. The quantities differ to no more than 3% between the data integration choices (a) and (b), with the brain stem being an exception, where the maximum difference between (a) and (b) is the fluid inflow rate from 6 to 24 h and it differs to 24 %. Overall, the results presented in Table S6 suggest that the quantities of interest computed from the identified velocity fields are robust with respect to data integration choices. This justifies using study averaged constants for $D$ and $T_1$ for the 9 subjects were this data is not available.

| ID | n | $\bar{\phi}$ (mm/h) | $\|\|c\|\|_0$ | $\|\|c_1\|\|_0$ | $\Delta c$ (%) | $\bar{\phi}$ (mm/h) | $\|\|c\|\|_0$ | $\|\|c_1\|\|_0$ | $\Delta c$ (%) | $\bar{\phi}$ (mm/h) | $\|\|c\|\|_0$ | $\|\|c_1\|\|_0$ | $\Delta c$ (%) |
|---|---|---|---|---|---|---|---|---|---|---|---|---|---|
| 3 | 16 | 0.27 | 50.01 | 45.40 | 10.16 | 0.51 | 46.14 | 45.40 | 1.63 | 0.60 | 45.06 | 45.40 | 0.76 |
| | 32 | 0.22 | 48.47 | 44.51 | 8.89 | 0.37 | 44.97 | 44.51 | 1.05 | 0.47 | 43.60 | 44.51 | 2.05 |
| 4 | 16 | 0.28 | 64.51 | 60.84 | 6.03 | 0.45 | 61.56 | 60.84 | 1.18 | 0.50 | 60.78 | 60.84 | 0.10 |
| | 32 | 0.24 | 64.68 | 61.36 | 5.41 | 0.32 | 61.76 | 61.36 | 0.65 | 0.33 | 61.37 | 61.36 | 0.03 |
| 7 | 16 | 0.20 | 53.26 | 50.82 | 4.80 | 0.34 | 51.31 | 50.82 | 0.95 | 0.40 | 50.95 | 50.82 | 0.26 |
| | 32 | 0.17 | 52.57 | 50.37 | 4.37 | 0.24 | 50.77 | 50.37 | 0.80 | 0.25 | 50.41 | 50.37 | 0.07 |
| 8 | 16 | 0.40 | 124.83 | 124.04 | 0.64 | 0.46 | 122.35 | 124.04 | 1.36 | 0.47 | 122.53 | 124.04 | 1.22 |
| | 32 | 0.28 | 125.82 | 125.25 | 0.45 | 0.31 | 123.62 | 125.25 | 1.30 | 0.31 | 123.68 | 125.25 | 1.25 |
| 9 | 16 | 0.14 | 71.20 | 70.04 | 1.67 | 0.18 | 70.14 | 70.04 | 0.16 | 0.19 | 69.98 | 70.04 | 0.08 |
| | 32 | 0.12 | 71.13 | 70.15 | 1.40 | 0.13 | 70.49 | 70.15 | 0.49 | 0.13 | 70.39 | 70.15 | 0.34 |
| 10 | 16 | 0.31 | 113.09 | 111.16 | 1.74 | 0.37 | 110.75 | 111.16 | 0.36 | 0.37 | 111.18 | 111.16 | 0.02 |
| | 32 | 0.22 | 114.14 | 112.87 | 1.12 | 0.25 | 112.69 | 112.87 | 0.16 | 0.25 | 112.39 | 112.87 | 0.42 |
| 11 | 16 | 0.23 | 112.90 | 111.50 | 1.26 | 0.29 | 111.40 | 111.50 | 0.09 | 0.29 | 111.39 | 111.50 | 0.10 |
| | 32 | 0.14 | 117.11 | 114.61 | 2.18 | 0.19 | 114.83 | 114.61 | 0.19 | 0.19 | 115.04 | 114.61 | 0.37 |
| 13 | 16 | 0.15 | 83.39 | 82.41 | 1.19 | 0.18 | 82.59 | 82.41 | 0.22 | 0.19 | 82.23 | 82.41 | 0.22 |
| | 32 | 0.12 | 84.11 | 83.11 | 1.20 | 0.13 | 83.66 | 83.11 | 0.66 | 0.13 | 83.57 | 83.11 | 0.55 |
| 15 | 16 | 0.09 | 47.94 | 47.26 | 1.43 | 0.11 | 47.51 | 47.26 | 0.52 | 0.13 | 47.36 | 47.26 | 0.21 |
| | 32 | 0.06 | 48.25 | 47.53 | 1.52 | 0.08 | 47.94 | 47.53 | 0.86 | 0.08 | 47.79 | 47.53 | 0.53 |
| 16 | 16 | 0.33 | 126.57 | 125.02 | 1.24 | 0.40 | 124.44 | 125.02 | 0.46 | 0.40 | 124.48 | 125.02 | 0.43 |
| | 32 | 0.24 | 126.05 | 125.54 | 0.41 | 0.24 | 125.93 | 125.54 | 0.31 | 0.24 | 125.88 | 125.54 | 0.27 |
| 17 | 16 | 0.19 | 106.83 | 105.63 | 1.14 | 0.22 | 105.85 | 105.63 | 0.22 | 0.23 | 105.66 | 105.63 | 0.03 |
| | 32 | 0.14 | 107.15 | 106.22 | 0.88 | 0.15 | 106.63 | 106.22 | 0.39 | 0.15 | 106.35 | 106.22 | 0.13 |
| 18 | 16 | 0.39 | 207.24 | 204.36 | 1.41 | 0.58 | 204.98 | 204.36 | 0.30 | 0.70 | 204.43 | 204.36 | 0.04 |
| | 32 | 0.35 | 213.71 | 211.22 | 1.18 | 0.50 | 211.76 | 211.22 | 0.26 | 0.55 | 211.51 | 211.22 | 0.14 |
| 19 | 16 | 0.13 | 98.99 | 98.14 | 0.86 | 0.16 | 98.32 | 98.14 | 0.18 | 0.16 | 98.23 | 98.14 | 0.09 |
| | 32 | 0.10 | 98.85 | 98.08 | 0.79 | 0.11 | 98.19 | 98.08 | 0.11 | 0.11 | 98.01 | 98.08 | 0.07 |
| 20 | 16 | 0.19 | 80.11 | 78.49 | 2.06 | 0.26 | 78.68 | 78.49 | 0.24 | 0.27 | 78.40 | 78.49 | 0.12 |
| | 32 | 0.16 | 79.89 | 78.35 | 1.97 | 0.19 | 78.52 | 78.35 | 0.22 | 0.20 | 78.52 | 78.35 | 0.21 |
| 21 | 16 | 0.14 | 112.86 | 112.70 | 0.14 | 0.16 | 112.44 | 112.70 | 0.24 | 0.16 | 112.42 | 112.70 | 0.25 |
| | 32 | 0.11 | 115.01 | 114.93 | 0.07 | 0.11 | 114.72 | 114.93 | 0.18 | 0.11 | 114.73 | 114.93 | 0.18 |
| 22 | 16 | 0.17 | 73.05 | 71.75 | 1.81 | 0.22 | 71.99 | 71.75 | 0.33 | 0.24 | 71.60 | 71.75 | 0.21 |
| | 32 | 0.14 | 72.63 | 71.41 | 1.71 | 0.17 | 71.89 | 71.41 | 0.67 | 0.17 | 71.59 | 71.41 | 0.26 |
| 23 | 16 | 0.11 | 221.48 | 221.29 | 0.08 | 0.12 | 221.29 | 221.29 | 0.00 | 0.12 | 221.25 | 221.29 | 0.02 |
| | 32 | 0.08 | 222.88 | 222.57 | 0.14 | 0.09 | 222.66 | 222.57 | 0.04 | 0.09 | 222.76 | 222.57 | 0.09 |
| 24 | 16 | 0.15 | 171.21 | 170.65 | 0.33 | 0.16 | 170.90 | 170.65 | 0.14 | 0.16 | 170.76 | 170.65 | 0.06 |
| | 32 | 0.10 | 176.80 | 176.37 | 0.24 | 0.11 | 176.53 | 176.37 | 0.09 | 0.11 | 176.42 | 176.37 | 0.03 |

DRAFT

**Supplementary Table S4. Evaluation of numerical accuracy of the estimated velocity fields and associated concentration fields for low ($n = 16$) and standard ($n = 32$) resolution meshes, and three values of the regularization parameter $\beta \in \{10^{-3}, 10^{-4}, 10^{-5}\}$ (set of columns) for the time intervals 24–48h. $\bar{\phi}$ (mm/h) is the brain-wide average velocity magnitude, $||c||_0$ the $L^2$-norm of the associated estimated concentration field, $||c_1||_0$ the $L^2$-norm of the corresponding target concentration field at end of the time interval (here ~48h), while $\Delta c$ (in percent) gives the relative difference between the latter two values.**

| | ID | $v$ ($\mu$m/min) | $v_g$ | $v_w$ | $v_s$ |
|---|---|---|---|---|---|
| 0 | 1 | 1.41 | 1.42 | 1.34 | 1.57 |
| 1 | 2 | 1.79 | 2.11 | 1.43 | 2.24 |
| 2 | 4 | 2.38 | 2.52 | 2.19 | 2.69 |
| 3 | 5 | 1.22 | 1.27 | 1.18 | 1.14 |
| 4 | 6 | 3.24 | 3.40 | 3.01 | 4.33 |
| 5 | 7 | 3.24 | 3.45 | 2.86 | 5.06 |
| 6 | 8 | 2.81 | 3.08 | 2.47 | 2.77 |
| 7 | 9 | 3.27 | 3.44 | 3.02 | 3.41 |
| 8 | 10 | 1.78 | 1.85 | 1.69 | 1.78 |
| 9 | 11 | 2.16 | 2.34 | 1.90 | 3.04 |
| 10 | 12 | 2.91 | 3.12 | 2.61 | 3.54 |
| 11 | 13 | 3.08 | 3.33 | 2.68 | 4.92 |
| 12 | 14 | 2.63 | 2.88 | 2.33 | 4.01 |
| 13 | 15 | 4.06 | 4.29 | 3.71 | 5.94 |
| 14 | 16 | 1.89 | 1.88 | 1.89 | 2.14 |
| 15 | 17 | 1.62 | 1.79 | 1.43 | 1.69 |
| 16 | 18 | 1.37 | 1.27 | 1.48 | 1.54 |
| 17 | 19 | 2.05 | 2.14 | 1.91 | 2.75 |
| 18 | 20 | 2.45 | 2.57 | 2.26 | 3.84 |
| 19 | 21 | 1.89 | 2.09 | 1.61 | 2.83 |
| 20 | 22 | 2.18 | 2.29 | 2.04 | 2.53 |
| 21 | 24 | 1.60 | 1.73 | 1.43 | 1.52 |
| Ref. (n=16) | | $2.40 \pm 0.82$ | $2.54 \pm 0.89$ | $2.10 \pm 0.72$ | $2.98 \pm 1.34$ |
| Dep. (n=6) | | $2.11 \pm 0.57$ | $2.28 \pm 0.60$ | $1.89 \pm 0.50$ | $2.92 \pm 1.29$ |
| Total (n=22) | | $2.32 \pm 0.75$ | $2.48 \pm 0.81$ | $2.11 \pm 0.67$ | $2.97 \pm 1.30$ |

**(a) 6–24h**

| | ID | $v$ ($\mu$m/min) | $v_g$ | $v_w$ | $v_s$ |
|---|---|---|---|---|---|
| 0 | 3 | 6.14 | 7.10 | 5.07 | 4.42 |
| 1 | 4 | 5.29 | 5.93 | 4.54 | 4.61 |
| 2 | 7 | 3.97 | 4.35 | 3.42 | 3.97 |
| 3 | 8 | 5.19 | 6.06 | 4.18 | 3.32 |
| 4 | 9 | 2.21 | 2.52 | 1.79 | 1.57 |
| 5 | 10 | 4.15 | 4.55 | 3.74 | 1.99 |
| 6 | 11 | 3.21 | 3.47 | 2.90 | 2.51 |
| 7 | 13 | 2.12 | 2.20 | 2.05 | 1.09 |
| 8 | 15 | 1.25 | 1.37 | 1.13 | 0.86 |
| 9 | 16 | 3.97 | 4.30 | 3.62 | 3.11 |
| 10 | 17 | 2.48 | 2.64 | 2.32 | 1.49 |
| 11 | 18 | 8.39 | 8.14 | 8.59 | 11.42 |
| 12 | 19 | 1.81 | 1.84 | 1.78 | 1.58 |
| 13 | 20 | 3.23 | 3.58 | 2.81 | 4.13 |
| 14 | 21 | 1.87 | 2.03 | 1.68 | 2.06 |
| 15 | 22 | 2.76 | 3.01 | 2.48 | 2.06 |
| 16 | 23 | 1.46 | 1.47 | 1.43 | 1.68 |
| 17 | 24 | 1.79 | 1.90 | 1.68 | 1.30 |
| Ref. (n=11) | | $4.23 \pm 1.98$ | $4.62 \pm 2.03$ | $3.77 \pm 1.98$ | $3.62 \pm 2.85$ |
| Dep. (n=7) | | $2.11 \pm 0.59$ | $2.24 \pm 0.69$ | $1.96 \pm 0.47$ | $1.90 \pm 1.03$ |
| Total (n=18) | | $3.41 \pm 1.89$ | $3.69 \pm 2.01$ | $3.07 \pm 1.79$ | $2.95 \pm 2.43$ |

**(b) 24–48h**

**Supplementary Table S5. Subject-specific average velocities $v$ denotes the brain-wide average flow speed (velocity magnitude, in $\mu$m/min), while $v_g$, $v_w$ and $v_s$ denote the cerebral cortex average, subcortical white matter average and brain stem average flow speed, respectively. Note the use of $\mu$m/min here and in Results versus mm/h for Table SM3. For the reference group at 6–24 hours, the regional average flow speeds mutually differ (paired t-test, p-values for cerebral cortex-subcortical white matter: 0.00016, cerebral cortex-brain stem: 0.011, white-brain stem: 0.0018). For the sleep-deprived group at 6–24h, the regional average flow speeds mutually differ, except when comparing the cerebral cortex/brain stem (paired t-test, p-values for cerebral cortex-subcortical white matter: 0.0018, cerebral cortex-brain stem: 0.075, subcortical white matter-brain stem: 0.031). The brain-wide and regional flow speeds do not differ between groups at 6h (Welch's t-test, p-values: 0.34, 0.40, 0.24, 0.79). For the groups combined, at 6–24h, the regional average velocities mutually differ (paired t-test, p-values for cerebral cortex-subcortical white matter: 1.1e-06, cerebral cortex-brain stem: 0.0013, subcortical white matter-brain stem: 8.9e-05). For the reference group at 24–48h, the subcortical white matter and cerebral cortex average flow speeds differ (paired t-test, p-values for cerebral cortex–subcortical white matter: 0.0031, cerebral cortex–brain stem: 0.23, subcortical white matter–brain stem: 0.87). For the sleep-deprived group at 24–48h, the subcortical white matter and cerebral cortex average flow speeds differ (paired t-test, p-values for cerebral cortex–subcortical white matter: 0.037, cerebral cortex–brain stem: 0.46, subcortical white matter–brain stem: 0.93).**

DRAFT

**Supplementary Table S6. Properties of the velocity field $\phi$ obtained with inverse modelling for two different preprocessing methods (using DTI & $T_1$ map vs using study averaged constants for $D$ and $T_1$), averaged over $n = 15$ subjects where both DTI and $T_1$ map is available.**

| | time interval | 6-24 h | | 24-48 h | |
|---|---|---|---|---|---|
| | method | T1 & DTI | avg. T1 & avgD | T1 & DTI | avg. T1 & avgD |
| Flow speed $\overline{\phi}$ ($\mu$m/min) | brain | 2.43 | 2.50 | 3.18 | 3.26 |
| | Cerebral cortex | 2.58 | 2.66 | 3.38 | 3.48 |
| | subcortical white matter | 2.19 | 2.27 | 2.92 | 2.99 |
| | brainstem | 3.53 | 3.34 | 3.20 | 3.03 |
| Fluid influx rate $\frac{1}{\lvert\Omega\rvert}\int_\Omega \nabla\cdot\phi\,\mathrm{d}x\ (10^{-4}\mathrm{min}^{-1})$ | Brain-wide | 0.95 | 0.95 | 2.67 | 2.72 |
| | Cerebral cortex | 2.55 | 2.58 | 2.92 | 2.93 |
| | Subcortical white matter | -1.14 | -1.13 | 2.32 | 2.44 |
| | brain stem | 3.10 | 2.37 | 3.54 | 3.00 |

### S4. Additional information for the inverse identification of dispersion and reaction parameters

This section gives additional methodological details and results for the inverse estimation of dispersion-reaction parameters, i.e. we consider finding $\alpha$ and $r$ in the following special case of Eq. (1) with $\phi = 0$

$$\frac{\partial c}{\partial t} - \nabla\cdot\alpha D^{*}\nabla c + rc = 0 \quad \text{in } \Omega, \quad t > t_0 \qquad \text{[S11]}$$

such that $c$ solves the PDE and minimizes the mismatch to the MRI observations.

**S4.1. Implementation details for inverse identification of dispersion and reaction parameters.** In order to solve Eq. (1) numerically, the equation is discretized in time via the Crank-Nicolson method and in space via the finite element method with continuous linear Lagrange elements. We find that for most subjects, a time step of 10 min yields convergence in the estimated dispersion and clearance parameters (cf. Section S4.3), but in some subjects (cf. Table S9), we performed additional simulations with time step size 5 min to ensure that the algorithm had converged. To obtain good starting values for the optimization of $\alpha, r$ we perform a grid search with combinations $\alpha \in \{1, 2, 3, 4, 5\}$, $r \in \{0, 6.25{\times}10^{-6}, 1.25{\times}10^{-5}, 2.5{\times}10^{-5}, 3.54{\times}10^{-5}, 5{\times}10^{-5}, 7.07{\times}10^{-5}, 10^{-4}\}$ for every subject. We find $\alpha = 3, r = 10^{-5}\,\mathrm{s}^{-1}$ as parameters that generally lead to significantly better agreement between simulations and data, and hence start the PDE-constrained optimization with these values for all subjects.

**S4.2. Influence of DTI and T1 map on parameter identification.** We further assessed how sensitive the identified dispersion-reaction parameters $(\alpha, r)$ are with respect to data integration choices. Using (a) subject-patient specific diffusion tensors and $T_1$ maps and (b) study averaged constants for $D$ and $T_1$ for the 15 subjects were this data is available, we compare optimal parameters obtained from PDE constrained optimization with a time step of 10 min in Table S7. For $\alpha$, the variation between methods is 0.002 which is much smaller than the variation $\sim 1.1$ between subjects, as measured by the standard deviation. With respect to the local clearance parameter $r$, the methods differ to $3.4 \times 10^{-4}\mathrm{min}^{-1}$ which is still one order of magnitude smaller than the variation between subjects ($\sim 15 \times 10^{-4}\mathrm{min}^{-1}$). Overall, the similarity of subject-specific optimal parameters presented in Table S7 suggest that both $\alpha$ and $r$ are robust with respect with respect to method (a) and (b). This justifies using study averaged constants for $D$ and $T_1$ for the 9 subjects were this data is not available.

DRAFT

**Supplementary Table S7. Dependence of optimal dispersion-reaction parameters on data integration choices. Time step = 10 min.**

| method | avg. $T_1$ & avg. $D$ | filtered $T_1$ & DTI | avg. $T_1$ & avg. $D$ | filtered $T_1$ & DTI |
|---|---|---|---|---|
| subject ID | $\alpha$ | | $r$ ($10^{-4}$ min$^{-1}$) | |
| 4 | 3.372 | 3.423 | 22.856 | 23.691 |
| 7 | 2.996 | 3.070 | 25.835 | 35.239 |
| 9 | 3.858 | 3.451 | 34.087 | 34.434 |
| 10 | 2.142 | 1.855 | 23.660 | 23.623 |
| 11 | 4.546 | 4.956 | 57.514 | 60.000 |
| 12 | 5.234 | 4.784 | 58.365 | 60.000 |
| 13 | 4.076 | 3.555 | 40.029 | 41.421 |
| 14 | 4.565 | 4.223 | 35.228 | 40.018 |
| 15 | 4.364 | 4.044 | 54.684 | 59.334 |
| 17 | 2.514 | 2.512 | 30.045 | 29.701 |
| 18 | 1.497 | 1.299 | 15.533 | 17.026 |
| 20 | 3.736 | 3.876 | 10.817 | 11.009 |
| 21 | 3.000 | 4.722 | 12.491 | 26.952 |
| 22 | 2.500 | 2.472 | 26.911 | 28.840 |
| 23 | 1.120 | 1.305 | 13.942 | 22.025 |
| mean ± std | 3.301 ± 1.152 | 3.303 ± 1.162 | 30.800 ± 15.418 | 34.221 ± 14.955 |

**S4.3. Numerical verification .** In order to ensure convergence of the PDE-constrained optimization algorithm with respect to the time step size used in the forward simulation, we performed computations with time step size of 30, 20 and 10 min for every subject. The results are given in Table S8. While the best parameters change between time step size 30 and 10 min, for most patients, the optimal parameters $\alpha$ and $r$ differ to no more than 10% between time step sizes 20 and 10 min. We hence consider the algorithm to be converged in these subject, and report the results obtained with time step 10 min in the manuscript. In those subjects where this criterion was not met, we perform additional simulations with time step size 5 min to ensure that the algorithm has converged. We find that in these subjects, the resulting $\alpha$ and $r$ differ to no more than 10% between time step sizes 10 and 5 min. We thus consider the algorithm to be converged in these subjects, and use the results obtained with time step size 5 min in these cases. For subject ID 3, the optimization algorithm converges to the upper bound $\alpha = 10$, and we hence excluded this subject from the results reported in the main text.

It should be noted that optimization for one subject typically requires around 15, 23, 49, and 81 hours for time steps 30, 20, 10 and 5 min, respectively, on a supercomputer node with 12 CPU cores.

**S4.4. Subject-level optimal enhanced diffusion-local clearance values.** Table S9 summarizes the results from PDE-constrained optimization for the best enhanced diffusion-local clearance parameters $\alpha, r$ for all subjects using averaged diffusion coefficients and $T_1$ values. The best parameters reduce the relative L2 mismatch to the MRI observations about $24 \pm 9\,\%$ (from $32 \pm 5\,\%$ to $24 \pm 6\,\%$) compared to diffusion simulations.

**Supplementary Table S9. Optimal enhanced diffusion-local clearance parameters.**

| Subject ID | sleep group? | best $\alpha$ | best $r$ $(10^{-4}\text{min}^{-1})$ | Rel. $L^2$-error with $(\alpha = 1, r = 0)$ | Rel. $L^2$-error with best $(\alpha, r)$ | red. in rel $L^2$-error (%) | time steps required for convergence |
|---|---|---|---|---|---|---|---|
| 1 | True | 3.9 | 40 | 0.29 | 0.25 | 13 | 288 |
| 2 | True | 2.8 | 49 | 0.37 | 0.34 | 7 | 288 |
| 4 | True | 3.4 | 23 | 0.26 | 0.20 | 24 | 576 |
| 5 | True | 1.5 | 22 | 0.24 | 0.20 | 15 | 288 |
| 6 | True | 7.0 | 11 | 0.34 | 0.23 | 32 | 576 |
| 7 | True | 3.1 | 26 | 0.28 | 0.24 | 16 | 576 |
| 8 | True | 6.6 | 39 | 0.40 | 0.31 | 23 | 576 |
| 9 | True | 3.8 | 34 | 0.36 | 0.30 | 16 | 576 |
| 10 | True | 2.1 | 24 | 0.28 | 0.16 | 42 | 288 |
| 11 | True | 4.8 | 62 | 0.40 | 0.31 | 22 | 576 |
| 12 | True | 5.2 | 59 | 0.37 | 0.30 | 21 | 576 |
| 13 | False | 4.1 | 40 | 0.26 | 0.17 | 35 | 288 |
| 14 | True | 4.6 | 35 | 0.39 | 0.31 | 19 | 288 |
| 15 | True | 4.4 | 55 | 0.31 | 0.24 | 22 | 288 |
| 16 | True | 2.9 | 25 | 0.28 | 0.20 | 27 | 288 |
| 17 | False | 2.5 | 30 | 0.26 | 0.17 | 35 | 288 |
| 18 | True | 1.5 | 16 | 0.24 | 0.15 | 37 | 288 |
| 19 | False | 3.3 | 33 | 0.34 | 0.24 | 29 | 288 |
| 20 | False | 3.7 | 11 | 0.28 | 0.23 | 19 | 576 |
| 21 | False | 3.0 | 12 | 0.36 | 0.31 | 15 | 288 |
| 22 | True | 2.5 | 27 | 0.29 | 0.23 | 20 | 288 |
| 23 | False | 1.1 | 14 | 0.31 | 0.18 | 43 | 288 |
| 24 | False | 2.6 | 24 | 0.28 | 0.22 | 22 | 576 |

DRAFT

### References.

1. PK Eide, V Vinje, AH Pripp, KA Mardal, G Ringstad, Sleep deprivation impairs molecular clearance from the human brain. *Brain* **144**, 863–874 (2021).
2. M Rohrer, H Bauer, J Mintorovitch, M Requardt, HJ Weinmann, Comparison of magnetic properties of mri contrast media solutions at different magnetic field strengths. *Investig. radiology* **40**, 715–724 (2005).
3. LM Valnes, et al., Apparent diffusion coefficient estimates based on 24 hours tracer movement support glymphatic transport in human cerebral cortex. *Sci. reports* **10**, 1–12 (2020).
4. AJ Taylor, M Salerno, R Dharmakumar, M Jerosch-Herold, T1 mapping: basic techniques and clinical applications. *JACC: Cardiovasc. Imaging* **9**, 67–81 (2016).
5. C Destrieux, B Fischl, A Dale, E Halgren, Automatic parcellation of human cortical gyri and sulci using standard anatomical nomenclature. *Neuroimage* **53**, 1–15 (2010).
6. B Fischl, Freesurfer. *Neuroimage* **62**, 774–781 (2012).
7. KA Mardal, ME Rognes, TB Thompson, LM Valnes, Mathematical modeling of the human brain: From magnetic resonance images to finite element simulation (2022).

**Supplementary Table S8. Dependence of optimal parameters on time steps size. In most subjects, the optimal parameters change less than 10% between time steps 20 and 10 min, and we hence did not run computationally expensive simulations with a time step of 5 minutes in these subjects.**

| pat | dt (min) | $\alpha$ | $r$ ($10^{-4}$ min$^{-1}$) | $J_p$ | $J_0$ | $J_F$ |
|---|---|---|---|---|---|---|
| 1 | 30 | 3.8 | 40 | 0.29 | 0.29 | 0.25 |
| | 20 | 3.9 | 41 | 0.29 | 0.29 | 0.25 |
| | 10 | 3.9 | 40 | 0.29 | 0.29 | 0.25 |
| | 5 | | | | | |
| 2 | 30 | 2.8 | 50 | 0.37 | 0.39 | 0.35 |
| | 20 | 2.8 | 51 | 0.37 | 0.39 | 0.34 |
| | 10 | 2.8 | 49 | 0.37 | 0.39 | 0.34 |
| | 5 | | | | | |
| 3 | 30 | 1.2 | 8 | 0.81 | 0.83 | 0.80 |
| | 20 | 3.0 | 14 | 0.50 | 0.46 | 0.46 |
| | 10 | 10.0 | 44 | 0.43 | 0.38 | 0.35 |
| | 5 | 10.0 | 47 | 0.43 | 0.35 | 0.35 |
| 4 | 30 | 2.1 | 14 | 0.46 | 0.44 | 0.42 |
| | 20 | 3.0 | 20 | 0.31 | 0.27 | 0.25 |
| | 10 | 3.4 | 23 | 0.26 | 0.21 | 0.19 |
| | 5 | 3.4 | 23 | 0.26 | 0.21 | 0.20 |
| 5 | 30 | 1.5 | 23 | 0.24 | 0.32 | 0.20 |
| | 20 | 1.4 | 21 | 0.24 | 0.32 | 0.20 |
| | 10 | 1.5 | 22 | 0.24 | 0.32 | 0.20 |
| | 5 | | | | | |
| 6 | 30 | 4.2 | 7 | 0.53 | 0.46 | 0.46 |
| | 20 | 5.9 | 10 | 0.37 | 0.29 | 0.27 |
| | 10 | 7.1 | 12 | 0.35 | 0.26 | 0.23 |
| | 5 | 7.0 | 11 | 0.34 | 0.26 | 0.23 |
| 7 | 30 | 1.8 | 15 | 0.40 | 0.40 | 0.37 |
| | 20 | 2.7 | 23 | 0.30 | 0.27 | 0.25 |
| | 10 | 3.0 | 26 | 0.28 | 0.26 | 0.24 |
| | 5 | 3.0 | 26 | 0.28 | 0.26 | 0.24 |
| 8 | 30 | 3.6 | 19 | 0.46 | 0.40 | 0.39 |
| | 20 | 5.5 | 31 | 0.41 | 0.34 | 0.32 |
| | 10 | 6.4 | 37 | 0.40 | 0.33 | 0.31 |
| | 5 | 6.6 | 39 | 0.40 | 0.33 | 0.31 |
| 9 | 30 | 1.9 | 16 | 0.67 | 0.66 | 0.63 |
| | 20 | 3.4 | 28 | 0.41 | 0.38 | 0.36 |
| | 10 | 3.9 | 34 | 0.36 | 0.32 | 0.30 |
| | 5 | 3.8 | 33 | 0.36 | 0.32 | 0.30 |
| 10 | 30 | 2.2 | 24 | 0.28 | 0.25 | 0.16 |
| | 20 | 2.3 | 25 | 0.28 | 0.25 | 0.16 |
| | 10 | 2.1 | 24 | 0.28 | 0.25 | 0.16 |
| | 5 | | | | | |
| 11 | 30 | 2.4 | 30 | 0.52 | 0.51 | 0.45 |
| | 20 | 4.0 | 50 | 0.42 | 0.39 | 0.33 |
| | 10 | 4.5 | 58 | 0.40 | 0.37 | 0.31 |
| | 5 | 4.8 | 62 | 0.40 | 0.37 | 0.31 |
| 12 | 30 | 3.2 | 33 | 0.42 | 0.38 | 0.35 |
| | 20 | 4.3 | 46 | 0.38 | 0.33 | 0.30 |
| | 10 | 5.2 | 58 | 0.38 | 0.33 | 0.30 |
| | 5 | 5.2 | 59 | 0.37 | 0.33 | 0.30 |
| 13 | 30 | 3.6 | 36 | 0.25 | 0.21 | 0.17 |
| | 20 | 4.1 | 40 | 0.26 | 0.21 | 0.17 |
| | 10 | 4.1 | 40 | 0.26 | 0.21 | 0.17 |
| | 5 | | | | | |

**(a) ID< 200**

| pat | dt (min) | $\alpha$ | $r$ ($10^{-4}$ min$^{-1}$) | $J_p$ | $J_0$ | $J_F$ |
|---|---|---|---|---|---|---|
| 14 | 30 | 4.5 | 36 | 0.39 | 0.34 | 0.32 |
| | 20 | 4.7 | 36 | 0.39 | 0.33 | 0.31 |
| | 10 | 4.6 | 35 | 0.39 | 0.33 | 0.31 |
| | 5 | | | | | |
| 15 | 30 | 4.1 | 51 | 0.31 | 0.28 | 0.24 |
| | 20 | 4.5 | 57 | 0.32 | 0.28 | 0.24 |
| | 10 | 4.4 | 55 | 0.31 | 0.28 | 0.24 |
| | 5 | | | | | |
| 16 | 30 | 2.7 | 24 | 0.28 | 0.24 | 0.20 |
| | 20 | 2.7 | 23 | 0.27 | 0.24 | 0.20 |
| | 10 | 2.9 | 25 | 0.28 | 0.24 | 0.20 |
| | 5 | | | | | |
| 17 | 30 | 2.3 | 27 | 0.26 | 0.25 | 0.17 |
| | 20 | 2.5 | 30 | 0.26 | 0.25 | 0.17 |
| | 10 | 2.5 | 30 | 0.26 | 0.25 | 0.17 |
| | 5 | | | | | |
| 18 | 30 | 1.5 | 16 | 0.24 | 0.25 | 0.15 |
| | 20 | 1.5 | 16 | 0.24 | 0.26 | 0.15 |
| | 10 | 1.5 | 16 | 0.24 | 0.26 | 0.15 |
| | 5 | | | | | |
| 19 | 30 | 2.9 | 30 | 0.34 | 0.30 | 0.24 |
| | 20 | 3.2 | 33 | 0.34 | 0.30 | 0.24 |
| | 10 | 3.3 | 33 | 0.34 | 0.30 | 0.24 |
| | 5 | | | | | |
| 20 | 30 | 3.0 | 9 | 0.28 | 0.23 | 0.23 |
| | 20 | 3.0 | 9 | 0.28 | 0.23 | 0.23 |
| | 10 | 3.7 | 11 | 0.28 | 0.23 | 0.23 |
| | 5 | 3.7 | 11 | 0.28 | 0.23 | 0.23 |
| 21 | 30 | 3.0 | 12 | 0.36 | 0.31 | 0.31 |
| | 20 | 3.0 | 12 | 0.36 | 0.32 | 0.31 |
| | 10 | 3.0 | 12 | 0.36 | 0.31 | 0.31 |
| | 5 | | | | | |
| 22 | 30 | 2.9 | 33 | 0.29 | 0.28 | 0.23 |
| | 20 | 2.6 | 28 | 0.29 | 0.28 | 0.23 |
| | 10 | 2.5 | 27 | 0.29 | 0.28 | 0.23 |
| | 5 | | | | | |
| 23 | 30 | 1.1 | 14 | 0.31 | 0.34 | 0.18 |
| | 20 | 1.1 | 14 | 0.31 | 0.33 | 0.17 |
| | 10 | 1.1 | 14 | 0.31 | 0.33 | 0.18 |
| | 5 | | | | | |
| 24 | 30 | 2.7 | 25 | 0.28 | 0.27 | 0.22 |
| | 20 | 3.0 | 29 | 0.28 | 0.28 | 0.22 |
| | 10 | 2.5 | 24 | 0.28 | 0.28 | 0.22 |
| | 5 | 2.6 | 24 | 0.28 | 0.27 | 0.22 |

**(b) ID > 200**

DRAFT
